# Structural basis for APE1 processing DNA damage in the nucleosome

**DOI:** 10.1101/2022.03.09.483662

**Authors:** Tyler M. Weaver, Nicole M. Hoitsma, Jonah J. Spencer, Lokesh Gakhar, Nicholas J. Schnicker, Bret D. Freudenthal

## Abstract

Genomic DNA is continually exposed to endogenous and exogenous factors that promote DNA damage. Eukaryotic genomic DNA is packaged into nucleosomes, which present a barrier to accessing and effectively repairing DNA damage. The mechanisms by which DNA repair proteins overcome this barrier to repair DNA damage in the nucleosome and protect genomic stability is unknown. Here, we determine how the base excision repair (BER) endonuclease AP-endonuclease 1 (APE1) recognizes and cleaves DNA damage in the nucleosome. Kinetic assays determined that APE1 cleaves solvent-exposed AP sites in the nucleosome with 3 - 6 orders of magnitude higher efficiency than occluded AP sites. A cryo-electron microscopy structure of APE1 bound to a nucleosome containing a solvent-exposed AP site identified that APE1 uses a DNA sculpting mechanism for AP site recognition, where APE1 bends the nucleosomal DNA to access the AP site. Notably, additional biochemical and structural characterization of occluded AP sites identified contacts between the nucleosomal DNA and histone octamer that prevent efficient processing of the AP site by APE1. These findings provide a rationale for the position-dependent activity of BER proteins in the nucleosome and suggests the ability of BER proteins to sculpt nucleosomal DNA drives efficient BER in chromatin.

## Main

Genomic DNA of eukaryotic cells is packaged into chromatin through a fundamental repeating unit called the nucleosome. The nucleosome consists of ~147bp of DNA wrapped around a core octameric histone protein complex, containing two copies each of histone H2A, H2B, H3, and H4^1^. The compact structure of the nucleosome and the robust contacts of the histone octamer with the nucleosomal DNA generates a barrier to accessing the genomic DNA sequence. Importantly, this nucleosome barrier must be overcome during critical cellular processes such as transcription, DNA replication, and DNA repair.

Both chromatinized and non-nucleosomal DNA are susceptible to endogenous and exogenous sources of DNA damage. One of the most common forms of DNA damage is the baseless sugar moiety known as apurinic/apyrimidinic (AP) sites, with an estimated ~10,000 AP sites generated in each cell per day^2,3^. These AP sites are generated through spontaneous depurination and depyrimidination, as well as through the excision of damaged DNA bases by damage-specific DNA glycosylases. Repair of AP sites is critical for maintaining genomic stability^4^ as they lack coding information during DNA replication^5^, are prone to generating cytotoxic DNA breaks^6^, and form DNA-protein crosslinks (DPCs) in the nucleosome^7,8^. The primary enzyme tasked with locating and processing genomic AP sites is the multifunctional nuclease AP endonuclease 1 (APE1), which is a key component of the base excision repair (BER) pathway^9^. During BER, APE1 functions as an AP-endonuclease by cleaving the DNA phosphodiester backbone 5′ of the AP site generating a 5′-nicked BER intermediate. The 5′-nicked BER intermediate is then processed by additional BER proteins, which ultimately restore the coding potential of the DNA to maintain genome stability. The biological importance of APE1, and the processing of genomic AP sites, is highlighted by the embryonic lethality of APE1 knockout in mice^10^, and the sensitivity of APE1-deficient cells to DNA damaging agents^11,12^.

The mechanisms used by APE1 and other BER enzymes to repair DNA damage in non-nucleosomal DNA are well-established^9,13,14^. However, how these enzymes function in the context of nucleosomes and higher-order chromatin remains poorly understood. *In-vitro* studies using recombinant nucleosomes have shown the activity of APE1 is highly dependent on the position of the DNA damage (i.e., the AP site) in the nucleosome, where solvent-exposed AP sites are more readily processed than occluded AP sites^15–18^. This position-dependent activity in the nucleosome is not exclusive to APE1 and is shared among other proteins in the BER pathway^18–27^. Consistent with these *in-vitro* observations, genome-wide repair of nucleosomal DNA base damage by BER proteins is also dependent on the position of the DNA damage in the nucleosome^28–30^. To date, the mechanisms BER proteins use for recognition and processing of DNA damage in the nucleosome, and the molecular basis for the position-dependent DNA repair activity remains unknown.

### APE1 AP-endonuclease activity is position-dependent in the context of the NCP

To investigate how the nucleosome impacts the AP-endonuclease activity of APE1, we generated three recombinant NCPs that each contain a single tetrahydrofuran AP site analogue. These three AP sites are positioned at distinct locations within the NCP described in terms of the position of the AP site relative to the nucleosome dyad, or superhelical location (SHL) (Fig, 1a and Extended Data Fig. 1a). Moreover, the AP sites also represent different rotational orientations, which dictates whether the phosphate backbone of the AP site is solvent-exposed (outward orientation) or near the core histone octamer (inward orientation). One nucleosome contains an AP site in a solvent-exposed rotational orientation near the nucleosome entry/exit site at SHL_−6_, referred to as AP-NCP_−6_. The second nucleosome contains an AP site in an inward facing rotational orientation near the nucleosome entry/exit site at SHL_−6.5_, referred to as AP-NCP_−6.5_. The third nucleosome contains an AP site in an inward facing rotational orientation one bp adjacent to the nucleosome dyad base at SHL_0_, referred to as AP-NCP_0_.

**Fig. 1.**
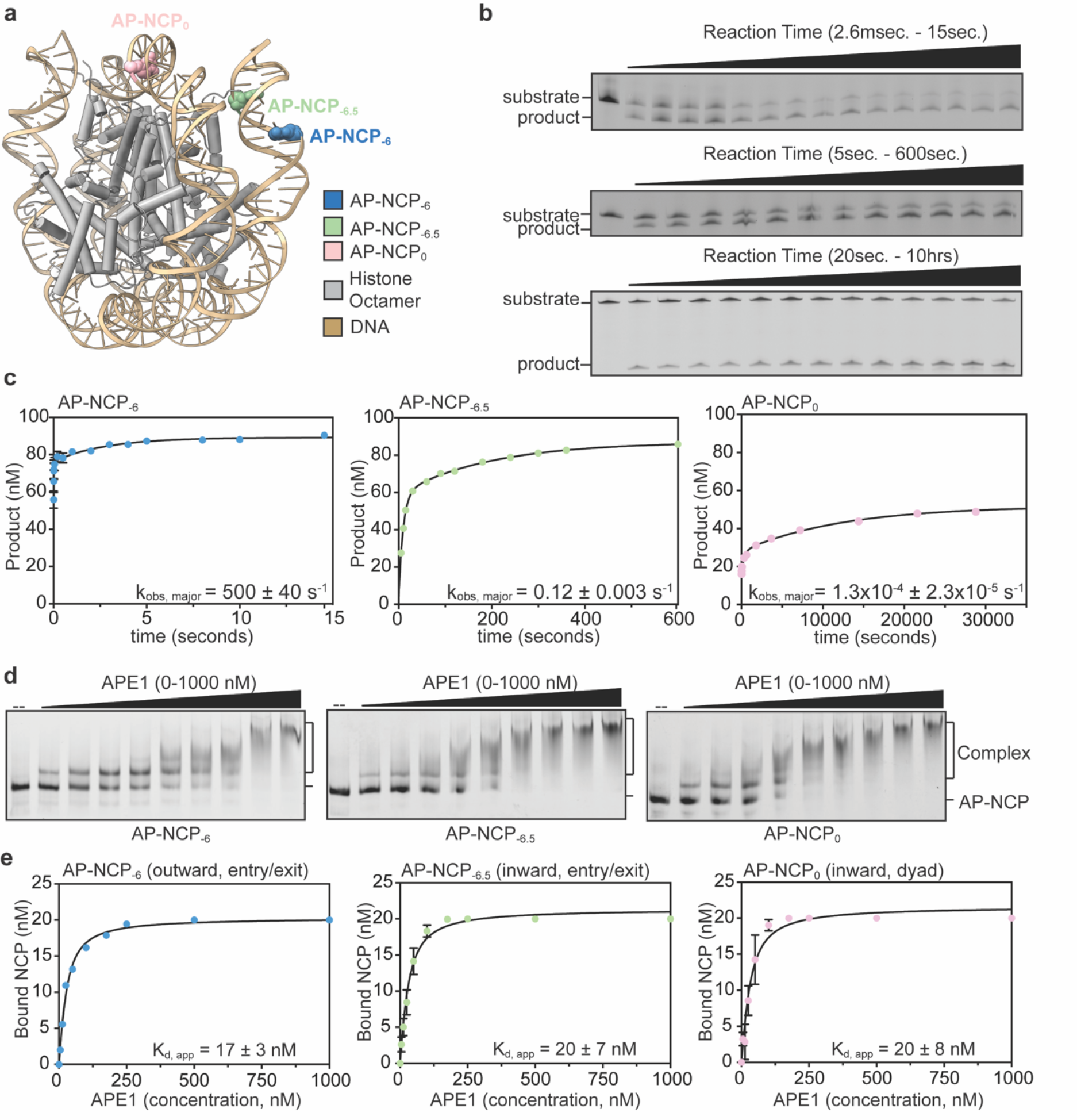
APE1 endonuclease activity is position-dependent in the nucleosome. **a,** Overall structure of a nucleosome core particle (PDB:4JJN) with the AP site positions at SHL_−6_, SHL_−6.5_, and SHL_0_ labeled. Representative gels for APE1 single turnover kinetic experiments (**b)** and quantification with fits **(c)** for AP-NCP_−6_, AP-NCP_−6.5_, and AP-NCP_0_. The data shown are the mean ± standard error of the mean from three replicate experiments. **d,** Representative APE1 EMSA gel **(d)** and quantification with fits **(e)** for AP-NCP_−6_, AP-NCP_−6.5_, and AP-NCP_0_. The data shown are the mean ± standard deviation from three replicate experiments. See Extended Data Fig. 1 for associated data.

To quantitatively characterize the position-dependent activity of APE1 on AP-NCPs, we performed single-turnover pre-steady state enzyme kinetics on AP-NCP_−6_, AP-NCP_−6.5_, and AP-NCP_0_ substrates (Fig. 1b and Extended Data Fig. 1c-d). Under single-turnover conditions, where APE1 is in excess of the AP-NCP substate, the kinetic time course corresponds to the first enzymatic turnover and can be fit to determine the APE1 DNA cleavage rate (k_obs_). As observed previously with non-nucleosomal DNA substrates^31^, APE1 exhibits biphasic kinetics with each of the AP-NCP substrates (Fig. 1c and Extended Data Fig. 1b). This kinetic behavior indicates the presence of two distinct cleavage rates. Here, we discuss the cleavage rate determined for the major population for each AP-NCP substrate as this likely represents the biologically relevant rate for AP-endonuclease activity. For AP-NCP_−6_, the observed rate constant for APE1 was 500 ± 40 s^−1^, which is similar to the k_obs_ of 441 ± 40 s^−1^ previously reported for APE1 on non-nucleosomal duplex DNA substrates. When the AP site was placed in a less solvent-exposed position near the nucleosome entry/exit site, AP-NCP_−6.5_, we observed a decrease in APE1 cleavage rate compared to the outward facing AP-NCP_−6_. The APE1 cleavage rate for AP-NCP_−6.5_ was 0.12 ± 0.01 s^−1^, which is a 3,650-fold decrease compared to non-nucleosomal duplex DNA. The largest reduction in APE1 activity was observed when the AP site was positioned in an inward facing orientation near the nucleosome dyad, AP-NCP_0_. The APE1 cleavage rate for AP-NCP_0_ was 1.30×10^−4^ ± 0.23×10^−4^ s^−1^, which is a 3,400,000-fold reduction compared to non-nucleosomal duplex DNA. To determine the specificity of APE1 cleavage for the nucleosomal AP sites, we generated a non-damaged nucleosome (ND-NCP) that lacks an AP site (Extended Data Fig. 1f). Importantly, APE1 cleavage activity was not observed for ND-NCPs, confirming the cleavage specificity of APE1 for nucleosomal AP sites (Extended Data Fig. 1g). Together, these pre-steady state kinetic measurements indicate that APE1 rapidly cleaves solvent-exposed AP sites at the nucleosome entry/exit site (AP-NCP_−6_). In contrast, less solvent-exposed AP sites at the nucleosome entry/exit (AP-NCP_−6.5_) and dyad (AP-NCP_0_) are cleaved with moderate and low efficiency, respectively.

To determine if the observed differences in APE1 cleavage observed at the three AP site positions is the result of reduced substrate binding, we performed electrophoretic mobility shift assays (EMSAs) with APE1 and AP-NCP_−6_, AP-NCP_−6.5_, and AP-NCP_0_ (Fig. 1d and Extended Data Fig. 1h-j). Quantification of the EMSAs yielded K_d,app_ of 17 ± 3 nM for AP-NCP_−6_, 20 ± 7 nM for AP-NCP_−6.5_, and 20 ± 8 nM for AP-NCP_0_ (Fig. 1e). Notably, the similar affinities of APE1 for all three AP-NCPs indicates the differences in catalytic efficiencies for AP-NCP_−6_, AP-NCP_−6.5_, and AP-NCP_0_ are not the result of reduced APE1 binding. To determine the contribution of the AP site to nucleosome binding, we performed additional EMSAs with ND-NCPs (Extended Data Fig. 3k). Quantification of the ND-NCP EMSAs yielded a K_d,app_ of 71 ± 10 nM, indicating APE1 has a ~3.5-fold higher affinity for AP-NCPs (Extended Data Fig. 3l).

### Mechanism for APE1 processing solvent-exposed AP sites in the nucleosome

The pre-steady state kinetics measurements indicate that APE1 rapidly processes a solvent-exposed AP site at SHL_−6_. To obtain mechanistic insight into how APE1 efficiently binds and processes the solvent-exposed AP site at SHL_−6_, we generated and purified an APE1-AP-NCP_−6_ substrate complex for structure determination by cryo-EM (Extended Data Fig. 2a). A subset of 58,854 particles was used to generate a 3.4 Å cryo-EM reconstruction of the APE1-AP-NCP_−6_ complex (Fig. 2, Extended Data Fig. 2, Extended Data Table 1). Both APE1 and the AP-NCP_−6_ are well-resolved in the cryo-EM map, with local resolutions of ~4.5 - 6 Å for APE1 and ~3 - 4 Å for AP-NCP_−6_ (Extended Data Fig. 2c - d). Interestingly, only the catalytic core of APE1 (residues 43 - 318) was observed in the APE1-AP-NCP_−6_ reconstruction despite utilizing full-length APE1 during cryo-EM grid preparation. The inability to resolve a large portion of the APE1 N-terminal domain (residues 1 - 42) in our cryo-EM reconstruction is likely due to significant conformationally flexibility and suggests it does not form a stable interaction with the nucleosome when bound at this AP site location. This conformational flexibility of the N-terminal domain is consistent with prior structural studies of APE1^32^.

**Fig. 2.**
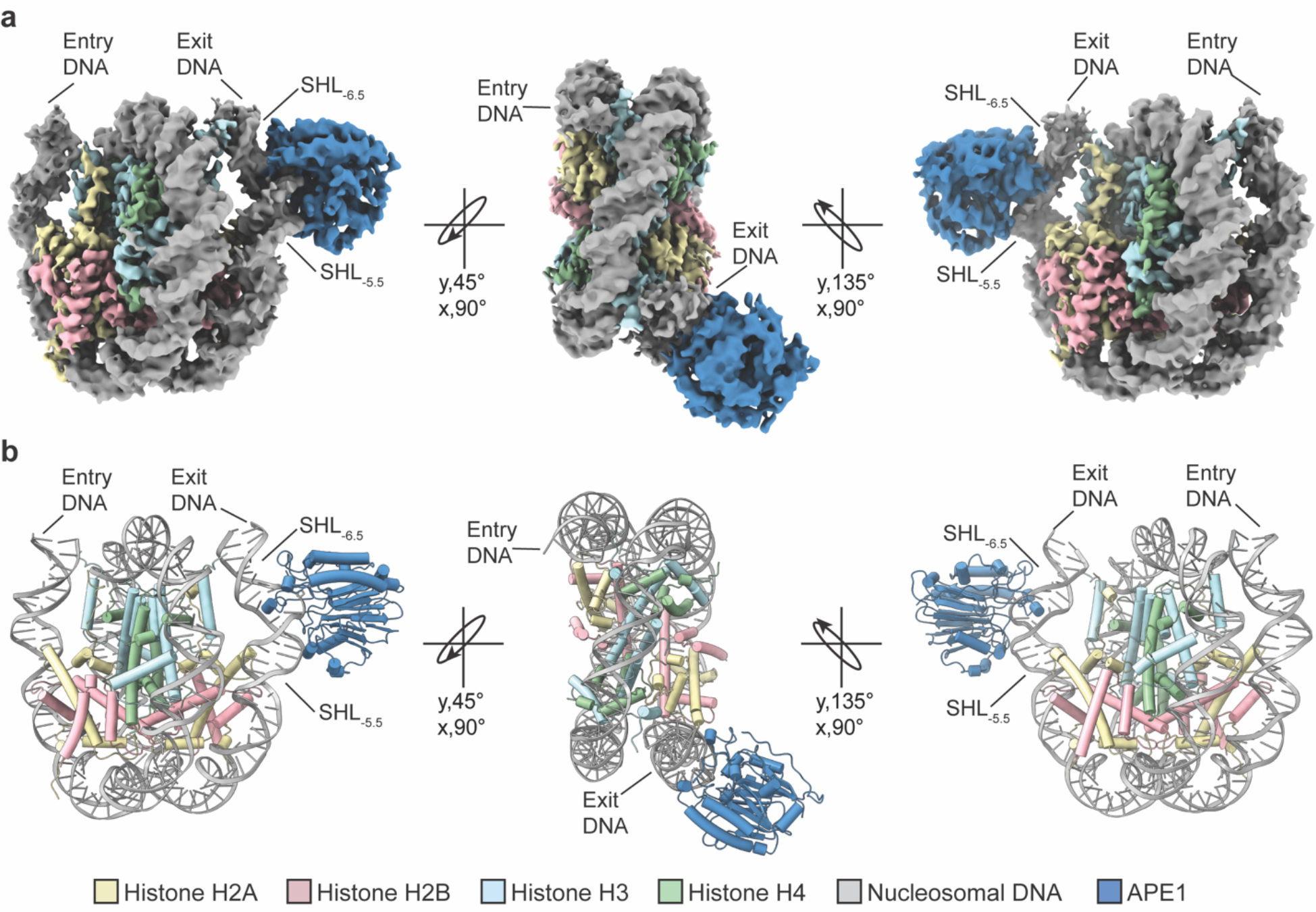
Cryo-EM structure of the APE1-AP-NCP_−6_ complex. **a,** A sharpened cryo-EM map of the APE1-AP-NCP complex in three different views. **b,** Model of the APE1- AP-NCP_−6_ complex shown in the same orientations as **(a)**.

In the APE1-AP-NCP_−6_ structure, APE1 is engaged with the nucleosome at the entry/exit site with a ~10 base-pair footprint that spans the nucleosomal DNA between SHL_−5.5_ to SHL_−6.5_ (Fig. 3a). Importantly, APE1 only interacts with the nucleosomal DNA and no direct contacts between APE1 and the core histone octamer are observed. The interaction between APE1 and the nucleosome occurs through three distinct molecular interfaces that make extensive contacts with both the non-damaged and damaged strands of the nucleosomal DNA (Fig. 3b). The first interface consists of the positively charged APE1 side chains of K103 and R73 that coordinate the phosphate backbone of the non-damaged DNA strand at SHL_−5.5_. (Fig. 3c). The second interface consists of a series of positively charged APE1 lysine side chains (K224, K227, K228) that are in position to interact with the negatively charged phosphate backbone of the non-damaged DNA strand near SHL_−6.5_ (Fig. 3d). The third interface is the APE1 active site, which interacts extensively with the nucleosomal DNA at SHL_−6_. The APE1 active site encompasses the nucleosomal DNA at SHL_−6_ by wedging R177 and the intercalating loop (residues 269 - 271) into the major and minor grooves of the nucleosomal DNA, respectively (Fig. 3e). In addition, the APE1 active site is in position to make extensive contacts with the damaged DNA strand surrounding the AP site (Fig. 3e, discussed below).

**Fig. 3.**
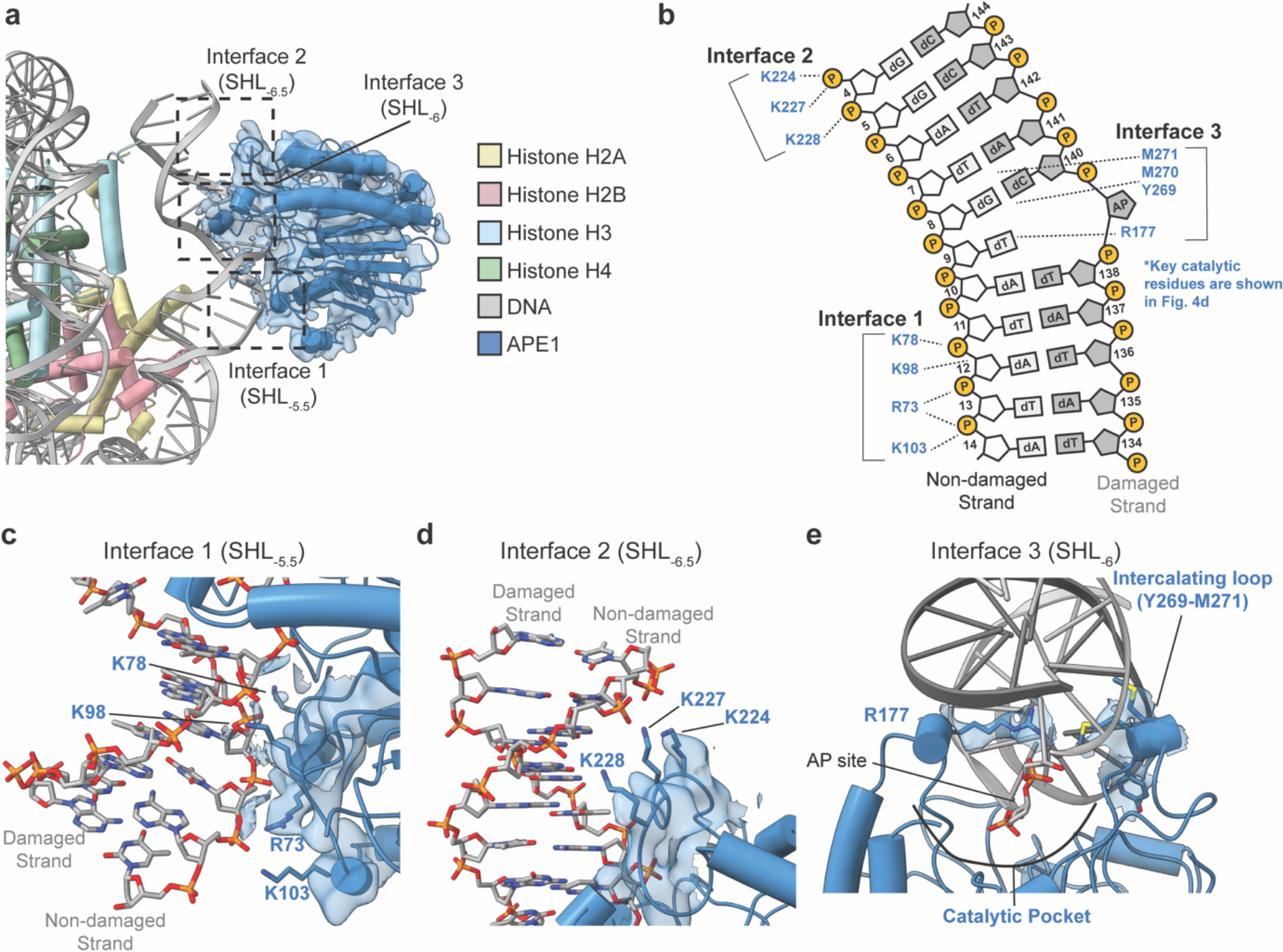
APE1 interacts with AP-NCP_−6_ using three molecular interfaces. **a,** A focused view of the APE1-AP-NCP_−6_ complex. Black dotted squares denote the three distinct molecular interfaces. The segmented APE1 cryo-EM map is shown as a transparent blue surface. **b,** A diagram representing the contacts between APE1 and the nucleosomal DNA identified using PLIP^66^. Several APE1 catalytic residues are not shown for clarity (see Fig. 4e). Focused view of **(c)** interface 1 at SHL_−5.5_, **(d)** interface 2 at SHL_−6.5_, **(e)** interface 3 at SHL_−6_ in the APE1-AP-NCP_−6_ complex. The cryo-EM map is shown as a transparent blue surface. Key APE1 active site residues are colored blue, but sticks representing some APE1 catalytic side chains are omitted for clarity in **(e)**.

APE1 utilizes a DNA sculpting mechanism for AP site recognition and catalysis in non-nucleosomal duplex DNA, where the DNA is bent ~35° to evict the AP site from the DNA helix into the APE1 active site^33,34^. To determine if APE1 similarly sculpts nucleosomal DNA, we obtained a 3.0 Å cryo-EM reconstruction of the AP-NCP_−6_ in the absence of APE1 (Extended Data Fig. 3 and Extended Data Table 1). Comparison of the APE1-AP-NCP_−6_ and AP-NCP_−6_ structures revealed significant distortion of the nucleosomal DNA upon APE1 binding (Fig. 4a). APE1 binding induces an additional ~20° bend in the nucleosomal DNA from SHL_−5.5_ to SHL_−6.5_ that results from a ~7 Å movement of the nucleosomal DNA away from the histone octamer at SHL_−6_ (Fig. 4a). In addition, APE1 binding results in significant widening of the minor groove of the nucleosomal DNA containing the AP site. The APE1-induced DNA bending does not cause significant structural rearrangements of the histone octamer, and contacts made between the histone octamer and the nucleosomal DNA at SHL_−5.5_ and SHL_−6.5_ remain intact (Extended Data Fig. 4a-c). In addition to DNA bending, significant variability in the APE1 position relative to the histone octamer was observed during 3D classification of the cryo-EM data (Extended Data Fig. 2b). Subsequent 3D variability analysis of the APE1-AP-NCP_−6_ particles revealed a substantial translational movement of APE1 around the nucleosomal DNA that is centered on the AP site, suggesting conformational heterogeneity during AP-site recognition in the nucleosome (Extended Data Fig. 4d).

**Fig. 4.**
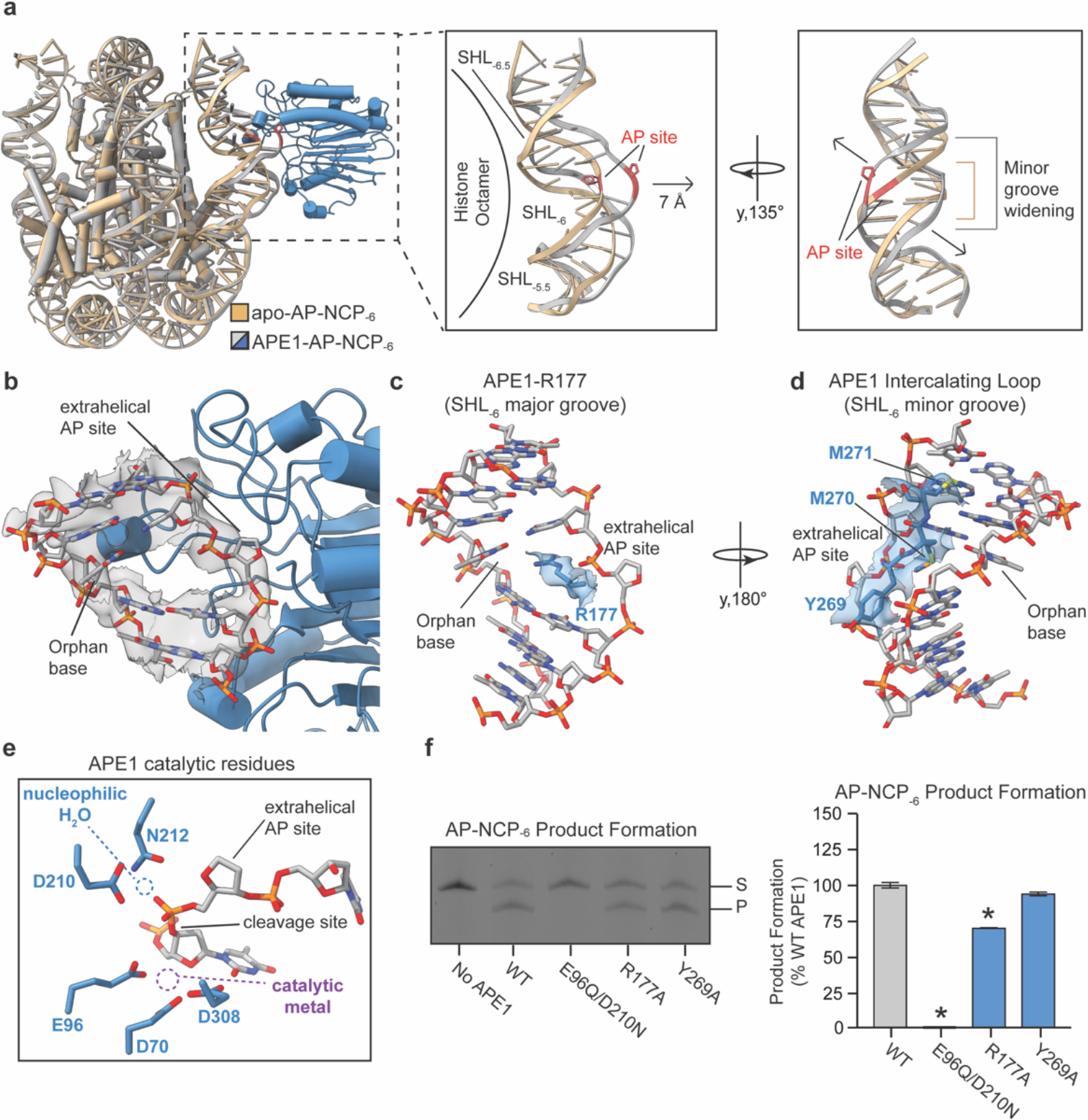
Fig. 4. Mechanism of nucleosomal AP site recognition by APE1. **a,** Structural overlay of the APE1-AP-NCP_−6_ and AP-NCP_−6_ structures. Focused view of the nucleosomal DNA between SHL_−5.5_ and SHL_−6.5_ without APE1 shown. **b,** Focused nucleosomal DNA view of the AP site in the APE1 active site. Cryo-EM map is shown as a transparent gray surface. **c-e,** Focused view of R177 in the APE1 active site at SHL_−6_ **(c)**, theAPE1 intercalating loop at SHL_−6_ **(d)**, and the nucleophilic water and metal coordinating residues in the APE1 active site **(e)**. The Cryo-EM map is shown as a transparent blue surface for **(d-e)**. The nucleophilic water and metal locations from PDB:5DFI are labeled in **(e)**. **f,** Representative gel (left) and quantification (right) of the AP-NCP_−6_ product formation assays for WT, E96Q/D210N, R177A, and Y269A APE1. The substrate (S) and product (P) are labeled. The data shown are the mean ± standard deviation from the three replicate experiments. The * denotes values that are significant different (p<0.01) than WT APE1 as determined by two-tailed student t-test.

The APE1-induced DNA bending facilitates displacement of the AP site from within the nucleosomal DNA helix into the APE1 active site (Fig. 4b). Importantly, this extrahelical conformation of the AP site is different than the predominantly intrahelical conformation seen in the AP-NCP_−6_ structure (Extended Data Fig. 4e). The void in the DNA helix generated by the extrahelical AP site is filled by R177, which wedges into the major groove of the nucleosomal DNA near SHL_−6_ (Fig. 4c). In this conformation, R177 sits across from the orphan DNA base of the non-damaged DNA strand and likely stabilizes the extrahelical conformation of the AP site. The APE1 active site makes additional contacts with the nucleosomal DNA at SHL_−6_ through intercalating loop residues Y269, M270, and M271, which wedge into the minor groove of the nucleosomal DNA (Fig. 4d). While several APE1 side chains were readily observed in the cryo-EM map, we did not observe clear density for all the side chains within the APE1 active site involved in catalysis. However, structural comparison of our APE1-AP-NCP_−6_ structure with a high-resolution crystal structure of APE1 bound to non-nucleosomal duplex DNA containing an AP site (APE1-AP-DNA, PDB:5DFI) revealed a similar mode of interaction with both substrates (Extended Data Fig. 4f)^33,34^. This enabled us to use the side-chain conformations from the high-resolution X-ray crystal structure for the remaining APE1 catalytic residues. Importantly, the positioning of the APE1 side chains that coordinate the nucleophilic water (residues D210 and N212) and catalytic metal (residues D70, E96, and D308) are in position to perform catalysis on the nucleosomal AP site (Fig 4e), suggesting APE1 uses a common catalytic mechanism for processing AP sites in the nucleosome and non-nucleosomal DNA

To obtain additional insight into the catalytic mechanism APE1 uses to cleave solvent-exposed AP sites in the nucleosome, we generated three APE1 active site variants (E96Q/D210N, R177A, and Y269A). In the APE1 AP-endonucleolytic cleavage reaction, residues E96 and D210 are critical for coordinating the catalytical metal and nucleophilic water^35–37^, respectively, R177 can act as a surrogate base to stabilize the extrahelical conformation of the AP site^33,38^, and Y269 is involved in APE1-mediated DNA sculpting^39^. We used a product formation assay to determine how well each APE1 mutant performs cleavage of AP-NCP_−6_ (Fig. 4f and Extended Data Fig. 5a). The APE1 E96Q/D210N mutant did not result in appreciable product formation for AP-NCP_−6_, suggesting this mutant is catalytically dead. The APE1 R177A mutant resulted in a significant decrease in product formation for AP-NCP_−6_, whereas Y269A had a minimal effect on product formation for AP-NCP_−6_ compared to WT APE1. This suggests that coordination of the catalytic metal and nucleophilic water, as well as stabilization of the extrahelical AP site conformation are important for APE1 cleavage of nucleosomal AP sites. Importantly, the E96Q/D210N, R177A, and Y269A mutants all maintain the ability to bind AP-NCP_−6_, indicating the reduction in product formation is not due to reduced nucleosome binding (Extended Data Fig. 5b). This structural and biochemical analysis are consistent with APE1 using a common mechanism for processing solvent-exposed AP sites in the nucleosome and AP sites in non-nucleosomal DNA. Importantly, the utilization of a common structural mechanism also provides a rationale for the similar cleavage rate of APE1 for AP-NCP_−6_ and AP-DNA (Fig. 1c and Extended Data Fig. 1b).

### Mechanism for APE1 processing occluded AP sites in the nucleosome

To further understand the mechanism APE1 uses to bind and cleave AP-NCP_−6.5_ and AP-NCP_0_, we attempted to determine structures of APE1 bound to AP-NCP_−6.5_ and AP-NCP_0_. Despite successfully generating APE1-AP-NCP_−6.5_ and APE1-AP-NCP_0_ complexes (Extended Data Fig. 6a and 7a), we were unable to obtain a reconstruction of APE1 bound to either AP-NCP. The exact reason for this is unclear, but likely indicates a high level of heterogeneity in APE1 binding position in the sample, which is consistent with the high affinity of APE1 to ND-NCPs (Extended Data Fig. 1k,l). To obtain additional insight into the catalytic mechanism APE1 uses to cleave APE1-AP-NCP_−6.5_ and APE1-AP-NCP_0_, we utilized the product formation assay to test the cleavage ability of the APE1 active site mutants (E98A/D210N, R177A, and Y269A) for AP-NCP_−6.5_ and AP-NCP_0_ (Fig. 5a and Extended Data Fig. 5c,e). The E96Q/D210N mutation completely abrogated APE1 endonuclease activity for AP-NCP_−6.5_ and AP-NCP_0_. In addition, the APE1 R177A and Y269A mutations resulted in a moderate decrease in product formation for AP-NCP_−6.5_ and AP-NCP_0_ compared to WT APE1. Importantly, the differences in product formation for AP-NCP_−6.5_ and AP-NCP_0_ are not due to reduced nucleosome binding (Extended Data Fig. 5d,f). The overall trends in the product formation assay for APE1 E96Q/D210A and R177A mutants cleaving AP-NCP_−6.5_ and AP-NCP_0_ are consistent with what was observed for AP-NCP_−6_, suggesting APE1 uses the same general catalytic mechanism for cleavage of all three AP-NCPs (compare Fig. 5a and 3d). Interestingly, the Y269A mutant had a larger effect on AP-NCP_−6.5_ and AP-NCP_0_ compared to the solvent-exposed AP-NCP_−6_, indicating Y269 may be important for DNA sculpting during attempts to access AP sites that are not solvent-exposed.

**Fig. 5.**
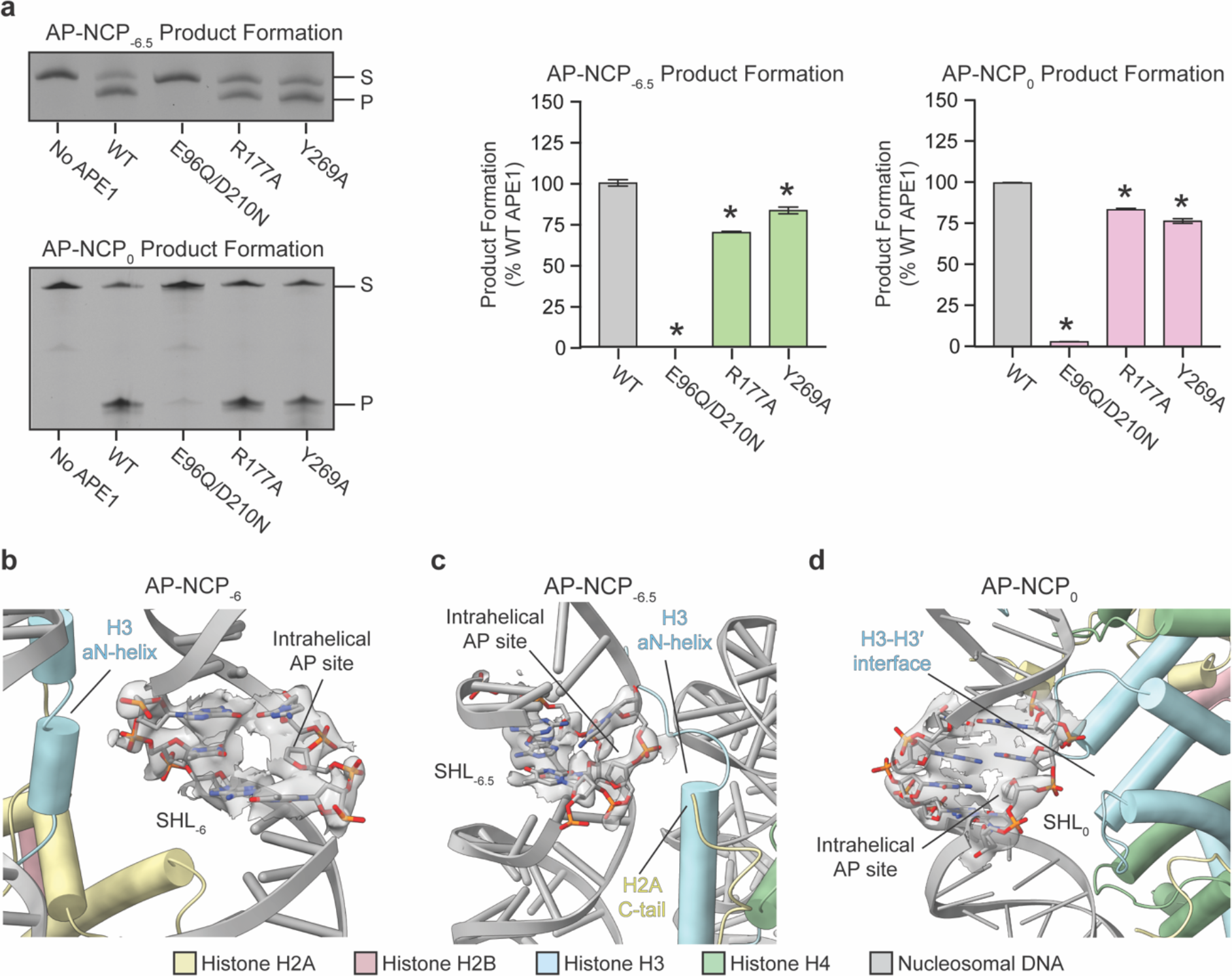
Mechanism for APE1 processing occluded AP sites in the nucleosome. **a,** Representative gels (left) and quantification (right) of the AP-NCP_−6.5_ and AP-NCP_0_ product formation assays for WT, E96Q/D210N, R177A, and Y269A APE1. The substrate (S) and product (P) are labeled. The data shown are the mean ± standard deviation from the three replicate experiments. The * denotes values that are significant different (p<0.01) than WT APE1 as determined by two-tailed student t-test. **b-d,** Focused views of the AP sites in AP-NCP_−6_ **(b)**, AP-NCP_−6.5_ **(c)**, and AP-NCP_0_ **(d)**. The cryo-EM map surrounding the AP site is shown as a transparent gray surface.

The product formation assays indicate APE1 uses a similar catalytic mechanism for cleaving all three AP-NCPs. This suggests the differences in APE1 cleavage rate for each AP-NCP may result from the inability of APE1 to access AP sites in the nucleosome. To determine any structural differences between the three AP-NCPs, we obtained two additional cryo-EM reconstructions of AP-NCP_−6.5_ and AP-NCP_0_ at 3.4 Å and 4.0 Å, respectively (Extended Data Fig. 6,7 and Extended Data Table 1). Structural comparison of the AP-NCP_−6_, AP-NCP_−6.5_, and AP-NCP_0_ revealed minimal changes to the overall structure of the nucleosome, indicating the three different AP sites do not induce large-scale conformational changes in the nucleosome (Extended Data Fig. 8a). In all three AP-NCP structures the AP site adopts a major conformation that is intrahelical, though we cannot rule out heterogeneity in the AP site conformation (Extended Data Fig. 8b). Closer inspection of three AP-NCP structures revealed significant differences in how extensive the nucleosomal DNA surrounding the AP sites interacts with the core histone octamer. The AP site at SHL_−6_ is completely solvent-exposed and no direct contacts between the histone octamer and damaged strand of the nucleosomal DNA are observed (Fig. 5b). In contrast, the damaged DNA strand containing the AP sites at SHL_−6.5_ and SHL_0_ make extensive interaction with the histone octamer. The AP site and surrounding nucleosomal DNA at SHL_−6.5_ interacts with the αN-helix of histone H3 and is in close proximity to the C-terminal tail of histone H2A (Fig. 5c). Similarly, the nucleosomal DNA surrounding the AP site at SHL_0_ is adjacent to the H3-H3′ interface at the nucleosome dyad (Fig. 5d). Subsequent modeling of APE1 bound at these occluded AP site positions revealed significant clashes between APE1 and the histone octamer that are incompatible with APE1 binding and DNA sculpting (Extended Data Fig. 8c). Together, this indicates large-scale structural rearrangements in the nucleosomal DNA and/or histone octamer are needed for processing occluded AP sites, which explains the moderate and low APE1 cleavage rates for AP-NCP_−6.5_ and AP-NCP_0_ (Fig. 1c and Extended Data Fig. 1b).

## Discussion

The cellular repair of AP sites is critical for maintaining genomic stability^4–6^. Our work describes how the essential BER protein APE1 recognizes and cleaves AP-sites in the nucleosome, providing a foundation for understanding how DNA repair occurs in chromatin. Our kinetic analysis revealed APE1 rapidly cleaves a solvent-exposed AP site near SHL_−6_, with a cleavage rate similar to non-nucleosomal DNA. The cryo-EM structure of the APE1- AP-NCP_−6_ complex provides a mechanistic basis for this observation. Nucleosomal AP site recognition by APE1 is accomplished through a DNA sculpting mechanism, where APE1 bends the DNA and flips the AP site out of the DNA helix and into its active site. Notably, the nucleosomal DNA sculpting by APE1 occurs without a direct interaction with histones or large structural rearrangements of the histone octamer. Similar nucleosomal DNA distortion was recently observed during nucleosome engagement by multiple pioneer transcription factors^40,41^, indicating this may be a common mechanism for accessing DNA within the nucleosome.

In contrast to the solvent-exposed AP site, the APE1 cleavage rates at occluded AP sites near SHL_−6.5_ and SHL_0_ are significantly lower, which is due to extensive interactions between the nucleosomal DNA and histone octamer that prevent efficient AP site binding, DNA sculpting, and catalysis by APE1. In this way, the histone octamer can serve as a barrier to DNA repair at the occluded AP site positions, which is consistent with genome-wide BER profiles^28–30^. While the details remain unknown, it is likely that both intrinsic nucleosome dynamics that enhance access to the DNA damage and/or extrinsic protein factors that alter the structure of the nucleosome are required for APE1 to overcome this barrier. At SHL_−6.5_, APE1 processes the AP site with a cleavage rate of ~0.1 s^−1^, which is almost identical to the equilibrium rate constant (0.02 - 0.1 s^−1^) for spontaneous unwrapping of the nucleosomal DNA at the entry/exit site^42^. Consistent with this finding, the histone PTM H3K56ac is known to increase spontaneous unwrapping of the nucleosomal DNA and was previously shown to enhance APE1 cleavage rate at an occluded nucleosomal AP site^15^. This suggests that intrinsic nucleosome dynamics likely facilitate repair of AP sites at occluded positions near the nucleosome entry/exit through a site-exposure model.

The AP site near the nucleosome dyad (SHL_0_) requires more extensive changes in nucleosome structure for efficient DNA repair. The UV-damaged DNA binding (UV-DDB) damage sensor protein, a recently identified BER co-factor^43^, uses a register shifting mechanism to alter the conformation of DNA damage from occluded to solvent-exposed positions within the nucleosome^44^. Additional BER co-factors PARP1 and the scaffolding protein XRCC1 are also known to directly and/or indirectly regulate nucleosome structure^24,45,46^. Finally, a variety of ATP-dependent chromatin remodeling complexes have been implicated in regulating BER protein activity in-vitro and cellular repair of DNA base damage^30,47–51^. These BER co-factors and chromatin remodeling enzymes would facilitate access of APE1 to occluded DNA damage in the nucleosome and enhance BER. However, the complex interplay between core BER factors, regulatory BER co-factors, and chromatin remodelers during DNA repair in chromatin remains poorly understood.

In addition to APE1, multiple BER proteins including DNA glycosylases and DNA polymerase β bend or sculpt the DNA from ~30° to 90° during damage recognition within non-nucleosomal DNA^13,14^ Our observation that APE1 adopts a similar conformation during AP site recognition in nucleosomal DNA and non-nucleosomal DNA indicates that DNA sculpting is likely needed for efficient repair in the nucleosome^33,34^. Whether DNA glycosylases and DNA polymerase β also sculpt nucleosomal DNA during damage recognition remains unknown, but the use of a DNA sculpting mechanism similar to that observed for APE1 would explain their position-dependent activity in the nucleosome^18,20–22,26,52–55^. Future work will be needed to identify the structural basis for DNA damage recognition by other core BER factors and whether these enzymes use a unified DNA sculpting mechanism for processing DNA damage in the nucleosome.

## Materials and Methods

### Purification of full-length APE1

Codon optimized human wild-type full-length (FL) APE1 in a pet28a vector was purchased from GenScript. The E96A/D210N, R177A, Y269A APE1 mutants were generated using the QuikChange II site-directed mutagenesis kit (Agilent). All proteins were expressed in One Shot BL21(DE3) plysS E. coli cells (Invitrogen), grown at 37 °C to an OD_600_ = 0.6, and induced with 0.4 mM IPTG overnight at 20 °C. Cells were subsequently harvested and lysed via sonication on ice in a buffer containing 50 mM HEPES (pH 7.4), 50 mM NaCl, and a protease inhibitor cocktail (AEBSF, leupeptin, benzamidine, pepstatin A). The cell lysate was cleared for 1 hour at 24,242 × g. The supernatant containing APE1 was purified via a HiTrap Heparin HP (GE Health Sciences) equilibrated with 50 mM HEPES (pH 7.4) and 50 mM NaCl, and APE1 eluted off the column with a linear salt gradient with 50 mM HEPES (pH 7.4) and 1 M NaCl. The eluted APE1 was diluted to 50 mM NaCl and further purified by cation-exchange chromatography using a POROS HS column (GE Health Sciences). APE1 was eluted from the POROS HS column with a linear salt gradient to 1 M NaCl. APE1 protein was further purified by gel filtration on a HiPrep 16/60 Sephacryl S-200 HR (GE Health Sciences). The purified APE1 protein was concentrated to >20 mg/mL and stored long term at −80 °C. All APE1 concentrations were determined via UV– Vis spectroscopy using a NanoDrop One Spectrophotometer (Thermo Scientific).

### Preparation of Oligonucleotides

To generate DNA substrates containing the 601 strong positioning sequence and single AP sites, a ligation-based method was utilized^56^. The individual DNA oligonucleotides (oligos) were synthesized by Integrated DNA Technologies (Extended Data Table 2). The oligos were resuspended in a buffer containing 10 mM Tris-pH 8.0 and 1 mM EDTA and mixed at a 1:1 ratio. The oligos were annealed by heating to 90 °C for 5 minutes before stepwise cooling to 4 °C using a linear gradient (−1 °C/min). The annealed DNA was subsequently ligated with T4 DNA ligase (CapeBio or New England Biolabs) and products separated via denaturing urea polyacrylamide gel electrophoresis (10% 37.5:1 acrylamide:bis-acrylamide). The ligated DNA was extracted from the gel in a buffer containing 200 mM NaCl and 1 mM EDTA using freeze-thaw cycles (3x, −20 °C to 37 °C). The purified and ligated DNA was reannealed heating to 90 °C for 5 minutes before stepwise cooling to 4 °C using a linear gradient (−1 °C/min). All ligated oligonucleotides were stored long term at −20 °C.

### Purification of Recombinant Histones

All histones were transformed and expressed in BL21 (DE3) (New England Biolabs) or BL21-CodonPlus (Agilent). Cells were grown in M9 minimal media at 37 °C until an OD_600_ of 0.4 was reached. Histone expression was induced with 0.3 mM (Histone H4) or 0.4 mM (Histone H2A, H2B, and H3) IPTG for 3-4 hours at 37 °C. Cell pellets were stored at −80 °C. Histone purification was performed using a previously established protocol^57^. In short, histones were extracted from inclusion bodies under denaturing conditions. Following extraction, the histones were further purified using a combination of anion-exchange and cation-exchange chromatography. The purified histones were dialyzed into H_2_O, lyophilized, and stored long term at −20 °C.

### Refolding of H2A/H2B Dimers and H3/H4 Tetramers

To generate H2A/H2B dimers and H3/H4 tetramers, each individual histone was resuspended in a buffer containing 20 mM Tris (pH 7.5), 6 M Guanidinium-HCl, and 10 mM DTT. The appropriate histones were mixed 1:1 molar ratio and dialyzed three time against a high salt buffer containing 2 M NaCl, 20 mM Tris (pH-7.5), and 1 mM EDTA. The refolded H2A/H2B dimers and H3/H4 tetramers were purified over a Sephacryl S-200 HR (GE Health Sciences) using high salt buffer containing 2 M NaCl, 20 mM Tris (pH-7.5), and 1 mM EDTA. The fractions containing pure H2A/H2B dimer and H3/H4 tetramer were combined and stored long term in 50% glycerol at −20 °C.

### Preparation of Nucleosomes

All nucleosomes were generated via a modified salt-dialysis method^57^. In short, H2A/H2B dimer, H3/H4 tetramer, and DNA were mixed at a 2:1:1 ratio, respectively. Nucleosomes were reconstituted via step-wise dialysis from 2 M NaCl to 1.5 M NaCl, 1.0 M NaCl, 0.66 M NaCl, 0.50 M NaCl, 0.25 M NaCl, and 0.125 M NaCl over 24-36 hours. The reconstituted nucleosomes were heat shocked at 55 °C to obtain uniform DNA positioning prior to purification via ultracentrifugation through a 10-40% sucrose gradient. Nucleosome formation and purity was determined by running native polyacrylamide gel electrophoresis (5% 59:1 acrylamide:bis-acrylamide).

### APE1 AP-Endonuclease Activity Assays

Single-turnover reactions were initiated by mixing full length APE1 enzyme (1000 nM) and AP-NCP substrate (100 nM) solutions in a reaction buffer containing 50 mM HEPES (pH-7.5), 100 mM KCl, 5 mM MgCl2 and 0.1 mg/ml bovine serum albumin (BSA) at 37 °C. A rapid quench flow system (KinTek RQF-3) was utilized for single-turnover pre-steady-state experiments time courses for AP-NCP_−6_ (2.6 msec. – 15 sec.), while time courses for AP-NCP_−6.5_ (5 sec. – 600 sec.) and AP-NCP_0_ (20 sec. – 10 hrs.) were completed in a benchtop heat block at 37 °C. The reactions were subsequently quenched with 300 mM EDTA at each respective time point. For mutant APE1 product formation assays, only a single timepoint was taken at 0.1 sec. for AP-NCP_−6_, 180 sec. for AP-NCP_−6.5_, and 10 hr. for AP-NCP_0_. All quenched reactions were mixed in 1:1 v/v ratio with a loading dye containing 100 mM EDTA, 80% deionized formamide, 0.25 mg/ml bromophenol blue and 0.25 mg/ml xylene cyanol. The reactions were incubated at 95°C for 6 min and separated by 15% 29:1 denaturing polyacrylamide gel electrophoresis. The bands corresponding to substrate and product were visualized by the 6-FAM label using an Amersham Typhoon RGB imager. The substrate and product bands were quantified using ImageQuant, and the data were best fit to the double exponential equation: Product = A(1−e^−*k*1*t^) + B(1−e^−*k*2*t^). Each time point represents an average of at least three independent replicate experiments ± the standard error of the mean. The indicated product formation for each APE1 mutant is the average of at least three independent replicate experiments ± the standard deviation.

### Electrophoretic mobility shift assays

Samples for electrophoretic mobility shift assays (EMSAs) were generated by mixing 20 nM AP-NCP with increasing concentrations of FL APE1 protein (5-1000 nM) in a buffer containing 10mM HEPES (7.5), 25 mM NaCl, 1 mM EDTA, 1 mM DTT, and 0.1 mg/mL BSA. For APE1 mutant EMSAs, samples were generated by mixing 20 nM AP-NCP with increasing concentrations of mutant APE1 protein (0 nM, 25 nM, and 250 nM) in a buffer containing 10mM HEPES (pH-7.5), 25 mM NaCl, 1 mM EDTA, 1 mM DTT, and 0.1 mg/mL BSA. The EMSA samples were mixed with equal volume 10% sucrose loading dye and complexes separated by native polyacrylamide gel (5%, 59:1 acrylamide:bis-acrylamide) electrophoresis in a running buffer containing 0.2x TBE for 45 minutes at 4 °C. The bands corresponding to free NCP and bound complexes were visualized by the 6-FAM label using an Amersham Typhoon RGB imager. The intensity of the bands was quantified using ImageJ^58^, and the data fit to a one-site binding model accounting for ligand depletion:

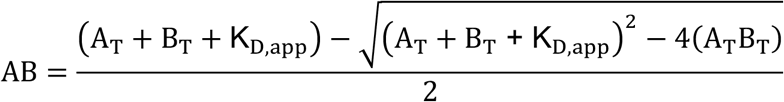

where A_T_ is the APE1 concentration, B_T_ is the NCP concentration, and AB is the concentration of APE1:NCP complex. Each data point represents an average of at least three independent replicate experiments ± the standard deviation. For the APE1 mutant EMSAs, three independent replicate experiments were performed.

### Cryo-EM sample and grid preparation

To generate APE1-AP-NCP_−6_, AP-NCP_−6_, AP-NCP_−6.5_, and AP-NCP_0_ samples, 125 μL of AP-NCP were incubated with 125 μL of FL APE1 at a 1:1.5 molar ratio (~6 μM AP-NCP and ~9 μM APE1) in a buffer containing 25 mM HEPES (pH 7.4), 50 mM NaCl, 1 mM TCEP, and 5 mM EDTA. The samples were incubated for 10 minutes at 4 °C before the addition of 0.20% glutaraldehyde for crosslinking. The samples were allowed to crosslink for an additional 10 minutes and immediately purified by gel filtration using a Superdex Increase 10/300 GL (GE Health Sciences) in a buffer containing 25 mM HEPES (pH 7.4), 50 mM NaCl, 1 mM TCEP, and 5 mM EDTA. Samples were concentrated to 0.20 mg/mL for APE1-AP-NCP_−6_, 0.20 mg/mL for AP-NCP_−6_, 0.25 mg/mL for AP-NCP_−6.5_, and 0.25 mg/mL for AP-NCP_0_ and subsequently stored at 4 °C until grid preparation. Gels corresponding to the samples used for cryo-EM grid preparation can be found in Extended Data Figs. 2a, 3a, 6a, 7a. Of note, the AP-NCP_−6_, AP-NCP_−6.5_, and AP-NCP_0_ samples were generated during attempts to crosslink APE1, though these attempts were unsuccessful and yielded only structures of the AP-NCPs (Extended Data Figs. 3, 6, 7).

Cryo-EM grids for the APE1-AP-NCP_−6_, AP-NCP_−6_, AP-NCP_−6.5_, and AP-NCP_0_ samples were generated by applying 3 μL (0.20-0.25 mg/mL) of each individual sample to a different Quantifoil R2/2 300 mesh copper grid that was glow-discharged for 60 seconds. The grids were blotted for 1-3 seconds at 8 °C and 95% humidity before being plunge-frozen in liquid ethane using an FEI Vitrobot Mark IV.

### Cryo-EM data collection and processing

All cryo-EM data collections were performed at the Pacific Northwest Center for Cryo-EM (PNCC) using an FEI Titan Krios 300 kV cryo-electron microscope equipped with a Gatan K3 Summit direct electron detector (Extended Data Table 1). For the APE1-AP-NCP_−6_ dataset, a total of 11,300 movies were recorded over ~48 hrs. from two separate cryo-EM grids. The data was collected using super-resolution mode with a pixel size of 0.40075 Å, a defocus range of −0.8 μm to −2.2 μm, and a total electron dose of 50 e^−^/Å^2^. For the AP-NCP_−6_ dataset, a total of 3,510 movies were recorded over ~24 hrs. from a single cryo-EM grid. The data was collected using super-resolution mode with a pixel size of 0.40075 Å, a defocus range of −0.8 μm to −2.2 μm, and a total electron dose of 50 e^−^/Å^2^. For the AP-NCP_−6.5_ dataset, a total of 3,680 movies were recorded over a ~24 hrs. from a single cryo-EM grid. The data was collected using super-resolution mode with a pixel size of 0.415 Å, a defocus range of −0.7 μm to −2.1 μm, and a total electron dose of 47 e^−^/Å^2^. For the AP-NCP_0_ dataset, a total of 4,995 movies were recorded over a single day from a single cryo-EM grid. The data was collected using super-resolution mode with a pixel size of 0.40075 Å, a defocus range of −0.8 μm to −2.0 μm, and a total electron dose of 50 e^−^/Å^2^.

All cryo-EM data processing was carried out using cryoSPARC^59^, and similar schemes were used to obtain each of the four cryo-EM structures. A correction for beam-induced drift was carried out using cryoSPARC patch motion correction and contrast transfer function (CTF) fit using cryoSPARC patch CTF-estimation. The micrographs were then manually curated. A random subset (~500) of the manually curated micrographs were used to perform block picking to generate templates for automated template picking. After template picking, a minimum of two rounds of 2D-classification were carried out to generate the final particle stacks. Ab-initio models were generated from the final particle stacks and multiple rounds of cryoSPARC heterogenous refinement were performed. The reconstructions for AP-NCP_−6_, AP-NCP_−6.5_, and AP-NCP_0_ were obtained after a final non-uniform refinement. After heterogenous refinement, a cryoSPARC 3D classification step was performed on the 294,348 particles of APE1-AP-NCP_−6_ complex using a mask for APE1 and the ~15bps of nucleosomal DNA composed of the APE1 binding site region. The class containing the most homogenous APE1-AP-NCP_−6_ complex was then subjected to a non-uniform refinement step to generate the final reconstruction. To resolve the conformational heterogeneity in the APE1-AP-NCP_−6_ complex, a three component cryoSPARC 3D variability analysis^60^ was performed on the initial 294,348 particles of the APE1-AP-NCP_−6_ complex. Further details for the cryo-EM processing pipeline for each of the four reconstructions can be found in Extended Data Figs. 2, 3, 6, 7.

The global resolution for all four structures was determined using Fourier shell correlation (FSC) 0.143 cur off. The AP-NCP_−6_, AP-NCP_−6.5_, and AP-NCP_0_ reconstructions were further subjected to B-factor sharpening using PHENIX^61^ autosharpen. For the APE1-AP-NCP_−6_ complex, the reconstruction was subjected to two separate B-factor sharpening steps. The first sharpening step was performed using the local resolution of the nucleosome and the second sharpening performed to the local resolution of the APE1/DNA region using PHENIX autosharpen^61^. The sharpened maps were then combined using PHENIX^61^ combine focused maps to yield the final composite map of the APE1-AP-NCP_−6_ complex.

### Model building and refinement

A human nucleosome model was generated using the coordinates from a yeast nucleosome structure (PDB:4JJN), which was chosen due to the strong similarity in the 601 positioning DNA sequence used in our study. The yeast histone residues were then mutated to the corresponding human histone residues using Coot^62^. The initial APE1 model was generated from a high-resolution X-ray crystal structure of an APE1-AP-DNA complex (PDB:5DFI)^33^. The initial models of the human nucleosome and APE1 were rigid-body docked into the APE1-AP-NCP_−6_ map using University of California San Francisco (UCSF) Chimera^63^. For AP-NCP_−6_, AP-NCP_−6.5_, and AP-NCP_0_, the initial model of the human nucleosome was rigid-body docked into the respective cryo-EM maps using UCSF Chimera^63^. All models were iteratively refined in PHENIX^61^ using secondary structure restraints and manual adjustments made to side chain conformations using Coot^62^. MolProbity^64^ was used to validate the final models prior to deposition. The model coordinates for the APE1-AP-NCP_−6_, AP-NCP_−6_, AP-NCP_−6.5_, and AP-NCP_0_ were deposited into the protein data bank under accession numbers 7U50, 7U51, 7U52, and 7U53. The cryo-EM maps for the APE1- AP-NCP_−6_, AP-NCP_−6_, AP-NCP_−6.5_, and AP-NCP_0_ were deposited into the electron microscopy data bank under accession numbers EMD-26336, EMD-26337, EMD-26338, and EMD-26339. All figures of the cryo-EM maps and models were generated using UCSF ChimeraX^65^.

## Acknowledgments

We thank Todd Washington (University of Iowa), Amy Whitaker (Fox Chase Cancer Center), and Alexandra Machen (University of Kansas Medical Center) for helpful discussion and assistance with the manuscript. We also thank Nancy Meyer for assistance with cryo-EM screening and data collection at PNCC. This research was supported by the National Institute of General Medical Science R35-GM128562 (B.D.F., T.M.W., N.M.H., and J.J.S.) and F32-GM140718 (T.M.W. and B.D.F.). A portion of this research was also supported by NIH grant U24GM129547 and performed at the PNCC at OHSU and accessed through EMSL (grid.436923.9), a DOE Office of Science User Facility sponsored by the Office of Biological and Environmental Research.

## Data Availability

Atomic coordinates for the reported structures have been deposited with the Protein Data bank under accession numbers 7U50, 7U51, 7U52, and 7U53. All cryo-EM maps are available from the electron microscopy data bank under accession numbers EMD-26336, EMD-26337, EMD-26338, and EMD-26339.

## References

1 Luger, K., Mäder, A. W., Richmond, R. K., Sargent, D. F. & Richmond, T. J. Crystal structure of the nucleosome core particle at 2.8 Å resolution. Nature 389, 251–260 (1997).

2 Lindahl, T. Instability and decay of the primary structure of DNA. nature 362, 709–715 (1993).

3 Nakamura, J. & Swenberg, J. A. Endogenous apurinic/apyrimidinic sites in genomic DNA of mammalian tissues. Cancer research 59, 2522–2526 (1999).

4 Loeb, L. A. & Preston, B. D. Mutagenesis by apurinic/apyrimidinic sites. Annual review of genetics 20, 201–230 (1986).

5 Cuniasse, P., Fazakerley, G., Guschlbauer, W., Kaplan, B. & Sowers, L. The abasic site as a challenge to DNA polymerase: a nuclear magnetic resonance study of G, C and T opposite a model abasic site. Journal of molecular biology 213, 303–314 (1990).

6 Kingma, P. S., Corbett, A. H., Burcham, P. C., Marnett, L. J. & Osheroff, N. Abasic Sites Stimulate Double-stranded DNA Cleavage Mediated by Topoisomerase II: DNA LESIONS AS ENDOGENOUS TOPOISOMERASE II POISONS (∗). Journal of Biological Chemistry 270, 21441–21444 (1995).

7 Zhou, C., Sczepanski, J. T. & Greenberg, M. M. Mechanistic studies on histone catalyzed cleavage of apyrimidinic/apurinic sites in nucleosome core particles. J Am Chem Soc 134, 16734–16741, doi:10.1021/ja306858m (2012).

8 Sczepanski, J. T., Zhou, C. & Greenberg, M. M. Nucleosome core particle-catalyzed strand scission at abasic sites. Biochemistry 52, 2157–2164, doi:10.1021/bi3010076 (2013).

9 Whitaker, A. M. & Freudenthal, B. D. APE1: A skilled nucleic acid surgeon. DNA repair 71, 93–100 (2018).

10 Xanthoudakis, S., Smeyne, R. J., Wallace, J. D. & Curran, T. The redox/DNA repair protein, Ref-1, is essential for early embryonic development in mice. Proceedings of the National Academy of Sciences 93, 8919–8923 (1996).

11 McNeill, D. R., Lam, W., DeWeese, T. L., Cheng, Y.-C. & Wilson, D. M. Impairment of APE1 function enhances cellular sensitivity to clinically relevant alkylators and antimetabolites. Molecular Cancer Research 7, 897–906 (2009).

12 Meira, L. B. et al. Heterozygosity for the mouse Apex gene results in phenotypes associated with oxidative stress. Cancer research 61, 5552–5557 (2001).

13 Brooks, S. C., Adhikary, S., Rubinson, E. H. & Eichman, B. F. Recent advances in the structural mechanisms of DNA glycosylases. Biochimica et Biophysica Acta (BBA)-Proteins and Proteomics 1834, 247–271 (2013).

14 Beard, W. A. & Wilson, S. H. Structure and mechanism of DNA polymerase β. Chemical reviews 106, 361–382 (2006).

15 Rodriguez, Y., Horton, J. K. & Wilson, S. H. Histone H3 Lysine 56 Acetylation Enhances AP Endonuclease 1-Mediated Repair of AP Sites in Nucleosome Core Particles. Biochemistry 58, 3646–3655, doi:10.1021/acs.biochem.9b00433 (2019).

16 Hinz, J. M. Impact of abasic site orientation within nucleosomes on human APE1 endonuclease activity. Mutation Research/Fundamental and Molecular Mechanisms of Mutagenesis 766–767, 19–24, doi:10.1016/j.mrfmmm.2014.05.008 (2014).

17 Hinz, J. M., Mao, P., McNeill, D. R. & Wilson, D. M., 3rd. Reduced Nuclease Activity of Apurinic/Apyrimidinic Endonuclease (APE1) Variants on Nucleosomes: IDENTIFICATION OF ACCESS RESIDUES. J Biol Chem 290, 21067–21075, doi:10.1074/jbc.M115.665547 (2015).

18 Beard, B. C., Wilson, S. H. & Smerdon, M. J. Suppressed catalytic activity of base excision repair enzymes on rotationally positioned uracil in nucleosomes. Proc Natl Acad Sci U S A 100, 7465–7470, doi:10.1073/pnas.1330328100 (2003).

19 Kennedy, E. E., Li, C. & Delaney, S. Global repair profile of human alkyladenine DNA glycosylase on nucleosomes reveals DNA packaging effects. ACS chemical biology 14, 1687–1692 (2019).

20 Bilotti, K., Tarantino, M. E. & Delaney, S. Human oxoguanine glycosylase 1 removes solution accessible 8-oxo-7, 8-dihydroguanine lesions from globally substituted nucleosomes except in the dyad region. Biochemistry 57, 1436–1439 (2018).

21 Cole, H. A., Tabor-Godwin, J. M. & Hayes, J. J. Uracil DNA glycosylase activity on nucleosomal DNA depends on rotational orientation of targets. Journal of Biological Chemistry 285, 2876–2885 (2010).

22 Tarantino, M. E., Dow, B. J., Drohat, A. C. & Delaney, S. Nucleosomes and the three glycosylases: High, medium, and low levels of excision by the uracil DNA glycosylase superfamily. DNA Repair (Amst) 72, 56–63, doi:10.1016/j.dnarep.2018.09.008 (2018).

23 Prasad, A., Wallace, S. S. & Pederson, D. S. Initiation of base excision repair of oxidative lesions in nucleosomes by the human, bifunctional DNA glycosylase NTH1. Mol Cell Biol 27, 8442–8453, doi:10.1128/MCB.00791-07 (2007).

24 Odell, I. D. et al. Nucleosome disruption by DNA ligase III-XRCC1 promotes efficient base excision repair. Mol Cell Biol 31, 4623–4632, doi:10.1128/MCB.05715-11 (2011).

25 Ye, Y. et al. Enzymatic excision of uracil residues in nucleosomes depends on the local DNA structure and dynamics. Biochemistry 51, 6028–6038, doi:10.1021/bi3006412 (2012).

26 Nilsen, H., Lindahl, T. & Verreault, A. DNA base excision repair of uracil residues in reconstituted nucleosome core particles. The EMBO journal 21, 5943–5952 (2002).

27 Rodriguez, Y. & Smerdon, M. J. The structural location of DNA lesions in nucleosome core particles determines accessibility by base excision repair enzymes. J Biol Chem 288, 13863–13875, doi:10.1074/jbc.M112.441444 (2013).

28 Pich, O. et al. Somatic and germline mutation periodicity follow the orientation of the DNA minor groove around nucleosomes. Cell 175, 1074–1087. e1018 (2018).

29 Mao, P. et al. Genome-wide maps of alkylation damage, repair, and mutagenesis in yeast reveal mechanisms of mutational heterogeneity. Genome research 27, 1674–1684 (2017).

30 Bohm, K. A. et al. Distinct roles for RSC and SWI/SNF chromatin remodelers in genomic excision repair. Genome research 31, 1047–1059 (2021).

31 Maher, R. L. & Bloom, L. B. Pre-steady-state kinetic characterization of the AP endonuclease activity of human AP endonuclease 1. Journal of Biological Chemistry 282, 30577–30585 (2007).

32 Tsutakawa, S. E. et al. Conserved structural chemistry for incision activity in structurally non-homologous apurinic/apyrimidinic endonuclease APE1 and endonuclease IV DNA repair enzymes. Journal of Biological Chemistry 288, 8445–8455 (2013).

33 Freudenthal, B. D., Beard, W. A., Cuneo, M. J., Dyrkheeva, N. S. & Wilson, S. H. Capturing snapshots of APE1 processing DNA damage. Nature structural & molecular biology 22, 924–931 (2015).

34 Mol, C. D., Izumi, T., Mitra, S. & Tainer, J. A. DNA-bound structures and mutants reveal abasic DNA binding by APE1 DNA repair and coordination. Nature 403, 451–456 (2000).

35 Nguyen, L. H., Barsky, D., Erzberger, J. P. & Wilson III, D. M. Mapping the protein-DNA interface and the metal-binding site of the major human apurinic/apyrimidinic endonuclease. Journal of molecular biology 298, 447–459 (2000).

36 McNeill, D. R. & Wilson, D. M. A dominant-negative form of the major human abasic endonuclease enhances cellular sensitivity to laboratory and clinical DNA-damaging agents. Molecular Cancer Research 5, 61–70 (2007).

37 Erzberger, J. P. & Wilson III, D. M. The role of Mg2+ and specific amino acid residues in the catalytic reaction of the major human abasic endonuclease: new insights from EDTA-resistant incision of acyclic abasic site analogs and site-directed mutagenesis. Journal of molecular biology 290, 447–457 (1999).

38 Liu, T.-C. et al. APE1 distinguishes DNA substrates in exonucleolytic cleavage by induced space-filling. Nature communications 12, 1–12 (2021).

39 Hoitsma, N. M. et al. AP-endonuclease 1 sculpts DNA through an anchoring tyrosine residue on the DNA intercalating loop. Nucleic acids research 48, 7345–7355 (2020).

40 Dodonova, S. O., Zhu, F., Dienemann, C., Taipale, J. & Cramer, P. Nucleosome-bound SOX2 and SOX11 structures elucidate pioneer factor function. Nature 580, 669–672 (2020).

41 Michael, A. K. et al. Mechanisms of OCT4-SOX2 motif readout on nucleosomes. Science 368, 1460–1465 (2020).

42 Li, G. & Widom, J. Nucleosomes facilitate their own invasion. Nature structural & molecular biology 11, 763–769 (2004).

43 Jang, S. et al. Damage sensor role of UV-DDB during base excision repair. Nature structural & molecular biology 26, 695–703 (2019).

44 Matsumoto, S. et al. DNA damage detection in nucleosomes involves DNA register shifting. Nature 571, 79–84 (2019).

45 Cannan, W. J., Rashid, I., Tomkinson, A. E., Wallace, S. S. & Pederson, D. S. The Human Ligase IIIalpha-XRCC1 Protein Complex Performs DNA Nick Repair after Transient Unwrapping of Nucleosomal DNA. J Biol Chem 292, 5227–5238, doi:10.1074/jbc.M116.736728 (2017).

46 van Beek, L. et al. PARP Power: A Structural Perspective on PARP1, PARP2, and PARP3 in DNA Damage Repair and Nucleosome Remodelling. International journal of molecular sciences 22, 5112 (2021).

47 Tsuda, M. et al. ALC1/CHD1L, a chromatin-remodeling enzyme, is required for efficient base excision repair. PLoS One 12, e0188320 (2017).

48 Verma, P. et al. ALC1 links chromatin accessibility to PARP inhibitor response in homologous recombination-deficient cells. Nature cell biology 23, 160–171 (2021).

49 Hewitt, G. et al. Defective ALC1 nucleosome remodeling confers PARPi sensitization and synthetic lethality with HRD. Molecular cell 81, 767–783. e711 (2021).

50 Charles Richard, J. L. et al. FACT Assists Base Excision Repair by Boosting the Remodeling Activity of RSC. PLoS Genet 12, e1006221, doi:10.1371/journal.pgen.1006221 (2016).

51 Menoni, H. et al. ATP-dependent chromatin remodeling is required for base excision repair in conventional but not in variant H2A.Bbd nucleosomes. Mol Cell Biol 27, 5949–5956, doi:10.1128/MCB.00376-07 (2007).

52 Hinz, J. M., Rodriguez, Y. & Smerdon, M. J. Rotational dynamics of DNA on the nucleosome surface markedly impact accessibility to a DNA repair enzyme. Proc Natl Acad Sci U S A 107, 4646–4651, doi:10.1073/pnas.0914443107 (2010).

53 Rodriguez, Y., Hinz, J. M., Laughery, M. F., Wyrick, J. J. & Smerdon, M. J. Site-specific Acetylation of Histone H3 Decreases Polymerase beta Activity on Nucleosome Core Particles in Vitro. J Biol Chem 291, 11434–11445, doi:10.1074/jbc.M116.725788 (2016).

54 Rodriguez, Y., Howard, M. J., Cuneo, M. J., Prasad, R. & Wilson, S. H. Unencumbered Pol beta lyase activity in nucleosome core particles. Nucleic Acids Res 45, 8901–8915, doi:10.1093/nar/gkx593 (2017).

55 Olmon, E. D. & Delaney, S. Differential Ability of Five DNA Glycosylases to Recognize and Repair Damage on Nucleosomal DNA. ACS Chem Biol 12, 692–701, doi:10.1021/acschembio.6b00921 (2017).

56 Bilotti, K., Kennedy, E. E., Li, C. & Delaney, S. Human OGG1 activity in nucleosomes is facilitated by transient unwrapping of DNA and is influenced by the local histone environment. DNA Repair (Amst) 59, 1–8, doi:10.1016/j.dnarep.2017.08.010 (2017).

57 Dyer, P. N. et al. Reconstitution of nucleosome core particles from recombinant histones and DNA. Methods in enzymology 375, 23–44 (2003).

58 Abràmoff, M. D., Magalhães, P. J. & Ram, S. J. Image processing with ImageJ. Biophotonics international 11, 36–42 (2004).

59 Punjani, A., Rubinstein, J. L., Fleet, D. J. & Brubaker, M. A. cryoSPARC: algorithms for rapid unsupervised cryo-EM structure determination. Nature methods 14, 290–296 (2017).

60 Punjani, A. & Fleet, D. J. 3D variability analysis: Resolving continuous flexibility and discrete heterogeneity from single particle cryo-EM. Journal of Structural Biology 213, 107702 (2021).

61 Adams, P. D. et al. PHENIX: a comprehensive Python-based system for macromolecular structure solution. Acta Crystallographica Section D: Biological Crystallography 66, 213–221 (2010).

62 Emsley, P. & Cowtan, K. Coot: model-building tools for molecular graphics. Acta crystallographica section D: biological crystallography 60, 2126–2132 (2004).

63 Pettersen, E. F. et al. UCSF Chimera—a visualization system for exploratory research and analysis. Journal of computational chemistry 25, 1605–1612 (2004).

64 Chen, V. B. et al. MolProbity: all-atom structure validation for macromolecular crystallography. Acta Crystallographica Section D: Biological Crystallography 66, 12–21 (2010).

65 Goddard, T. D. et al. UCSF ChimeraX: Meeting modern challenges in visualization and analysis. Protein Science 27, 14–25 (2018).

66 Adasme, M. F. et al. PLIP 2021: Expanding the scope of the protein–ligand interaction profiler to DNA and RNA. Nucleic acids research 49, W530–W534 (2021).

