## Extended data for "Structural basis for APE1 processing DNA damage in the nucleosome"

**Extended Data Fig. 1. Analysis of APE1 nucleosome binding and cleavage.**

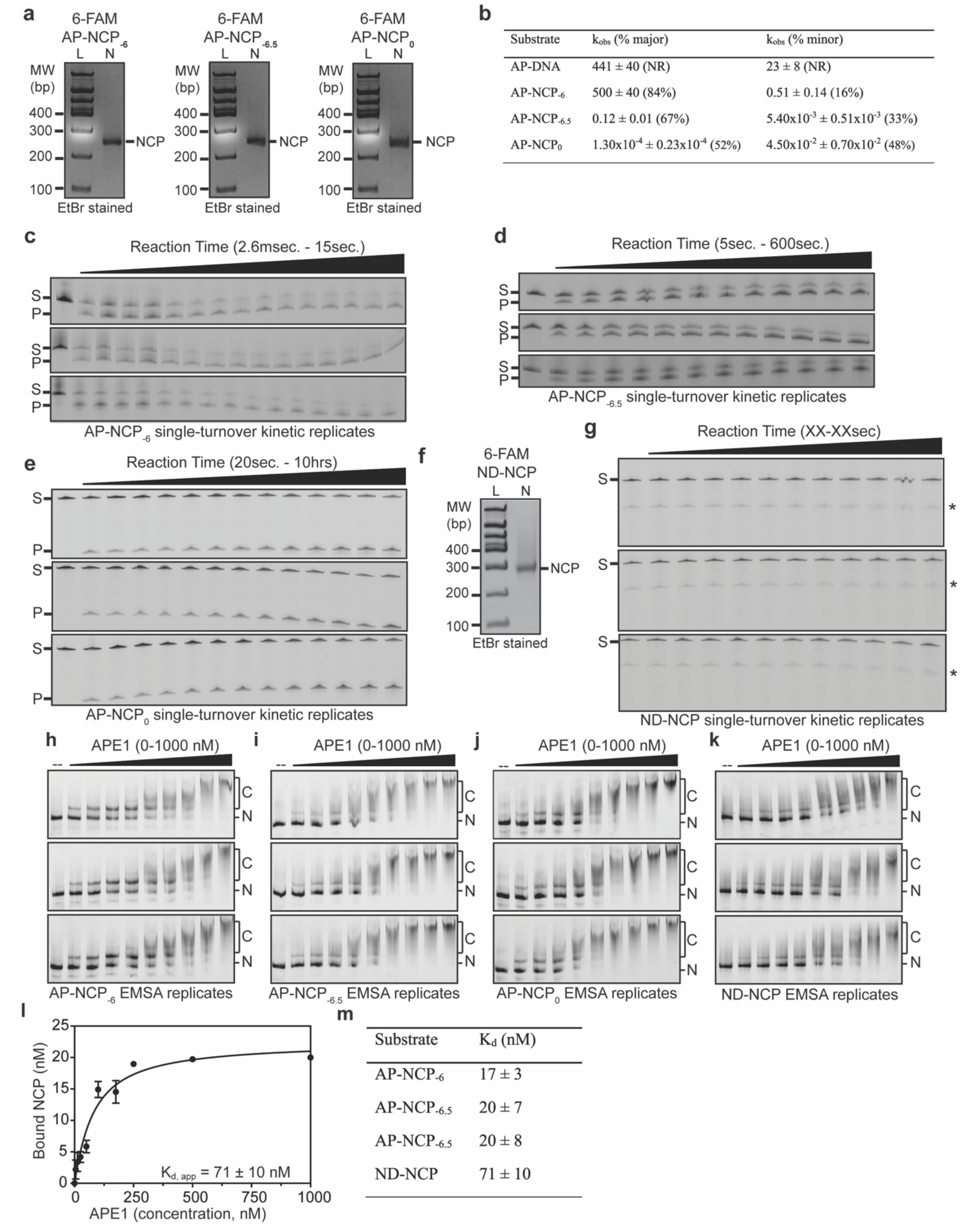


| Substrate | $k_{obs}$ (% major) | $k_{obs}$ (% minor) |
|---|---|---|
| AP-DNA | 441 ± 40 (NR) | 23 ± 8 (NR) |
| AP-NCP$_{-6}$ | 500 ± 40 (84%) | 0.51 ± 0.14 (16%) |
| AP-NCP$_{-6.5}$ | 0.12 ± 0.01 (67%) | $5.40x10^{-3} \pm 0.51x10^{-3}$ (33%) |
| AP-NCP$_{0}$ | $1.30x10^{-4} \pm 0.23x10^{-4}$ (52%) | $4.50x10^{-2} \pm 0.70x10^{-2}$ (48%) |

| Substrate | $K_d$ (nM) |
|---|---|
| AP-NCP$_{-6}$ | 17 ± 3 |
| AP-NCP$_{-6.5}$ | 20 ± 7 |
| AP-NCP$_{-6.5}$ | 20 ± 8 |
| ND-NCP | 71 ± 10 |

**Extended Data Fig. 1. Analysis of APE1 nucleosome binding and cleavage.**

**a,** Native PAGE gels confirming nucleosome formation for 6-FAM labeled AP-NCP$_{-6}$, AP-NCP$_{-6.5}$, and AP-NCP$_{0}$. The 100 bp DNA ladder (L) and nucleosome sample (S) are labeled. **b,** APE1 single turnover kinetic parameters for AP-NCP$_{-6}$, AP-NCP$_{-6.5}$, and AP-NCP$_{0}$. The kinetic parameters for AP-DNA were previously reported[31]. NR stands for not reported. **c-e**, Gels from three replicate APE1 single turnover pre-steady state kinetic experiments for AP-NCP$_{-6}$ **(c)**, AP-NCP$_{-6.5}$ **(d)**, and AP-NCP$_{0}$ **(e)**. The substrate (S) and product (P) are labeled. **f,** Native PAGE gels of 6-FAM labeled ND-NCP. The 100 bp DNA ladder (L) and nucleosome sample (S) are labeled. **g,** APE1 single turnover pre-steady state kinetic experiments for ND-NCPs. The substrate (S) is labeled. **h-k,** Gels from three replicate APE1 EMSA for AP-NCP$_{-6}$ **(h)**, AP-NCP$_{-6.5}$ **(i)**, AP-NCP$_{0}$ **(j)**, and ND-NCP **(k)**. The free nucleosome (N) and complex (C) are labeled **l,** Quantification of the ND-NCP EMSAs. The data shown is the mean ± standard deviation from the three replicate experiments. **m,** Table summarizing the $K_{d,app}$ calculated from the EMSAs for AP-NCP$_{-6}$, AP-NCP$_{-6.5}$, AP-NCP$_{0}$, and ND-NCP.

**Extended Data Fig. 2. Single particle analysis of the APE1-AP-NCP-6 complex.**

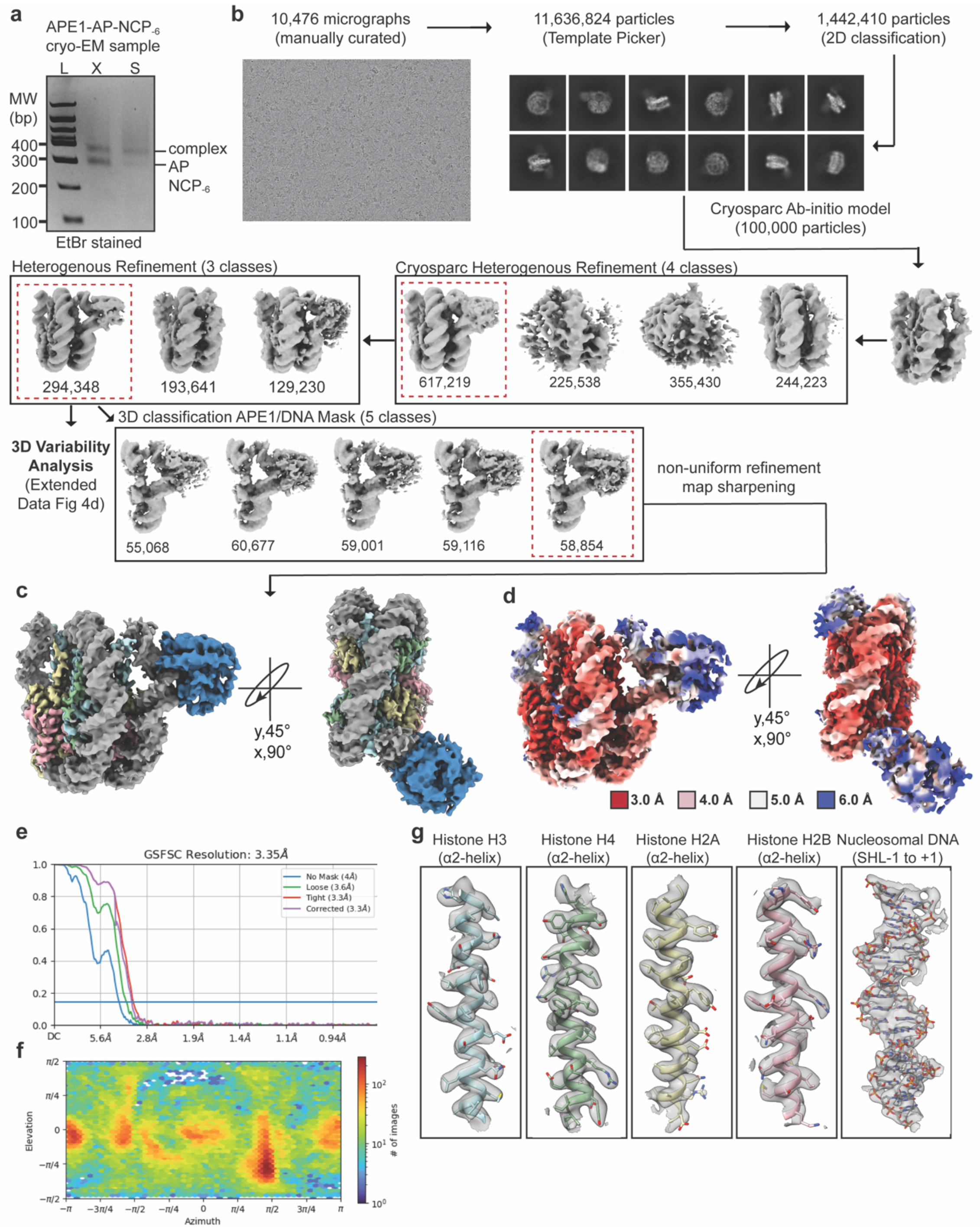

**Extended Data Fig. 2. Single particle analysis of the APE1-AP-NCP$_{-6}$ complex.**

**a,** Native PAGE gel of the APE1-AP-NCP$_{-6}$ cryo-EM sample. The 100 bp DNA ladder (L) and sample used to generate cryo-EM grids (S) are labeled. **b,** For single particle analysis, micrographs were manually curated and multiple rounds of 2D classification were performed yielding a final stack of 1,442,410 particles. A representative micrograph and set of 2D-classes are shown. An ab-initio model from 100,000 particles was generated prior to two rounds of heterogenous refinement (4 and 3 classes). A map containing APE1-AP-NCP$_{-6}$ was further subjected to 3D classification using a mask for APE1 and the nucleosomal DNA between SHL$_{-5.5}$ to SHL$_{-6.5}$. All maps chosen for downstream analysis are labeled by a dotted red box. **c,** Final 3.4 Å sharpened cryo-EM map of the APE1-AP-NCP$_{-6}$ complex. **d,** Local resolution estimate for the APE1-AP-NCP$_{-6}$ cryo-EM map. **e,** Fourier shell correlation (FSC-0.143) for the AP-NCP$_{-6}$ map. **f,** Heatmap of the angular distribution of particles used to generate the final APE1-AP-NCP$_{-6}$ cryo-EM map. **g,** Representative segmented density for H2A, H2B, H3, H4 and the nucleosomal DNA from the APE1-AP-NCP$_{-6}$ cryo-EM map.

**Extended Data Fig. 3. Single particle analysis of AP-NCP$_{-6}$.**

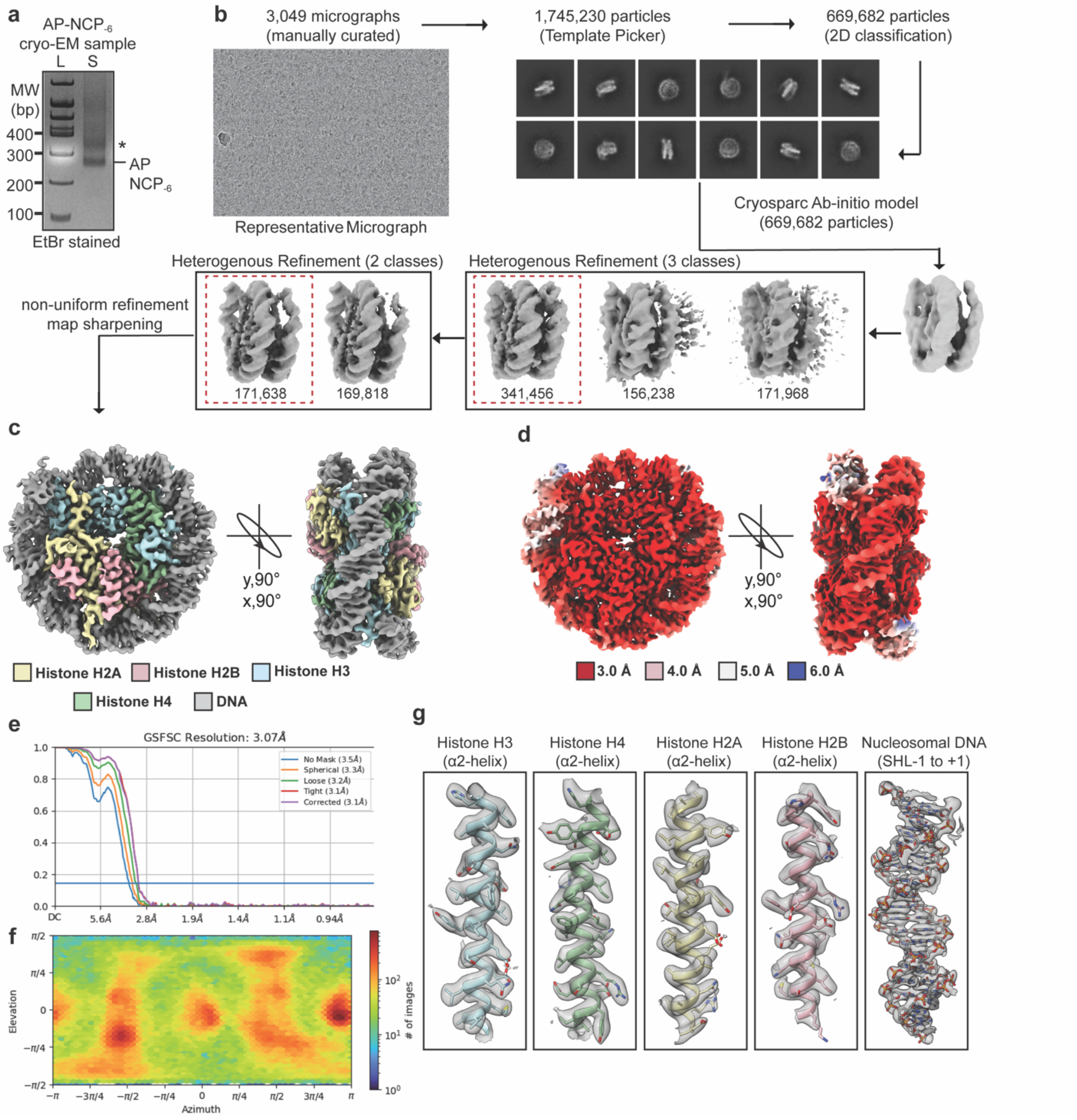

**Extended Data Fig. 3. Single particle analysis of AP-NCP$_{-6}$.**

**a,** Native PAGE gel of the purified AP-NCP$_{-6}$ cryo-EM sample. The 100 bp DNA ladder (L) and sample used to generate cryo-EM grids (S) are labeled. **b,** For single particle analysis, micrographs were manually curated and multiple rounds of 2D classification were performed yielding a final stack of 669,682 particles. A representative micrograph and set of 2D-classes are shown. An ab-initio model from 669,682 particles was generated before two rounds of heterogenous refinement (3 and 2 classes). All maps chosen for downstream analysis are labeled by a dotted red box. **c,** Final 3.1 Å sharpened cryo-EM map of AP-NCP$_{-6}$. **d,** Local resolution estimate for the AP-NCP$_{-6}$ cryo-EM map. **e,** Fourier shell correlation (FSC-0.143) for the AP-NCP$_{-6}$ map. **f,** Heatmap of the angular distribution of particles used to generate the final AP-NCP$_{-6}$ cryo-EM map. **g,** Representative segmented density for H2A, H2B, H3, H4 and the nucleosomal DNA from the AP-NCP$_{-6}$ cryo-EM map.

**Extended Data Fig. 4. Mechanism of nucleosomal AP site recognition by APE1.**

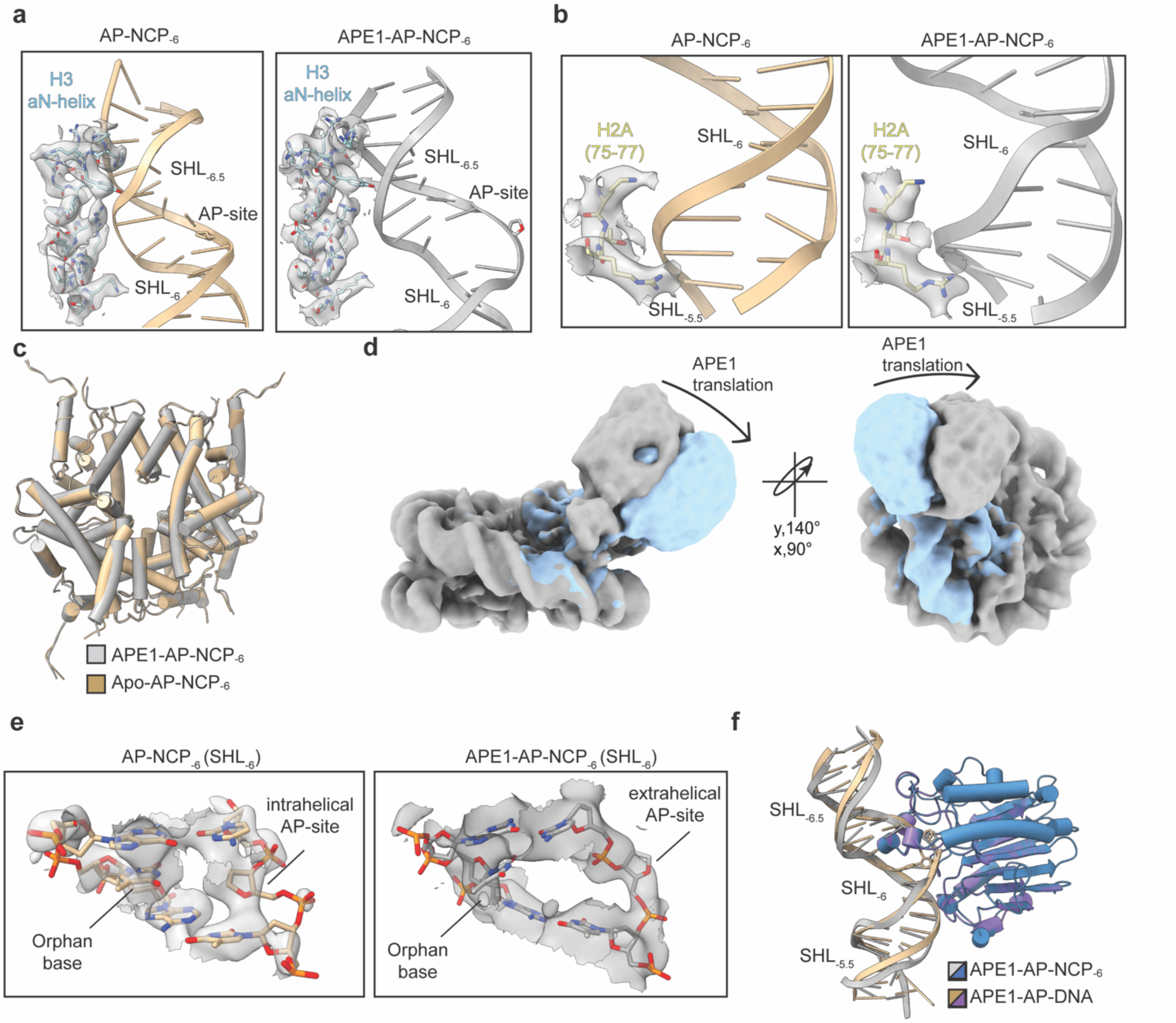

**Extended Data Fig. 4. Mechanism of nucleosomal AP site recognition by APE1.**

**a,** Focused views of the interaction between the H3 αN-helix and the nucleosomal DNA at $SHL_{-6.5}$ for $AP\text{-}NCP_{-6}$ (left) and the $APE1\text{-}AP\text{-}NCP_{-6}$ complex (right). The cryo-EM density for the H3 αN-helix is shown as a transparent gray surface. **b,** Focused views of the interactions between the H2A (residues 75-77) and the nucleosomal DNA at $SHL_{-5.5}$ for $AP\text{-}NCP_{-6}$ (left) and $APE1\text{-}AP\text{-}NCP_{-6}$ complex (right). Cryo-EM density for H2A (residues 75-77) is shown as a transparent gray surface. **c,** Structural comparison of the histone octamer from the $APE1\text{-}AP\text{-}NCP_{-6}$ and $AP\text{-}NCP_{-6}$ structures showing minimal structural rearrangements. **d,** Translational movement of APE1 around the nucleosomal DNA containing the AP site identified through 3D variability analysis. **e,** Focused views of the AP site at $SHL_{-6}$ for $AP\text{-}NCP_{-6}$ (left) and the $APE1\text{-}AP\text{-}NCP_{-6}$ complex (right). The cryo-EM density for the nucleosomal DNA is shown as a transparent gray surface. **f,** Structural comparison of the $APE1\text{-}AP\text{-}NCP_{-6}$ and APE1-AP-DNA (PDB:5DFI) complex structures.

**Extended Data Fig. 5. Analysis of nucleosome binding and cleavage by APE1 mutants.**

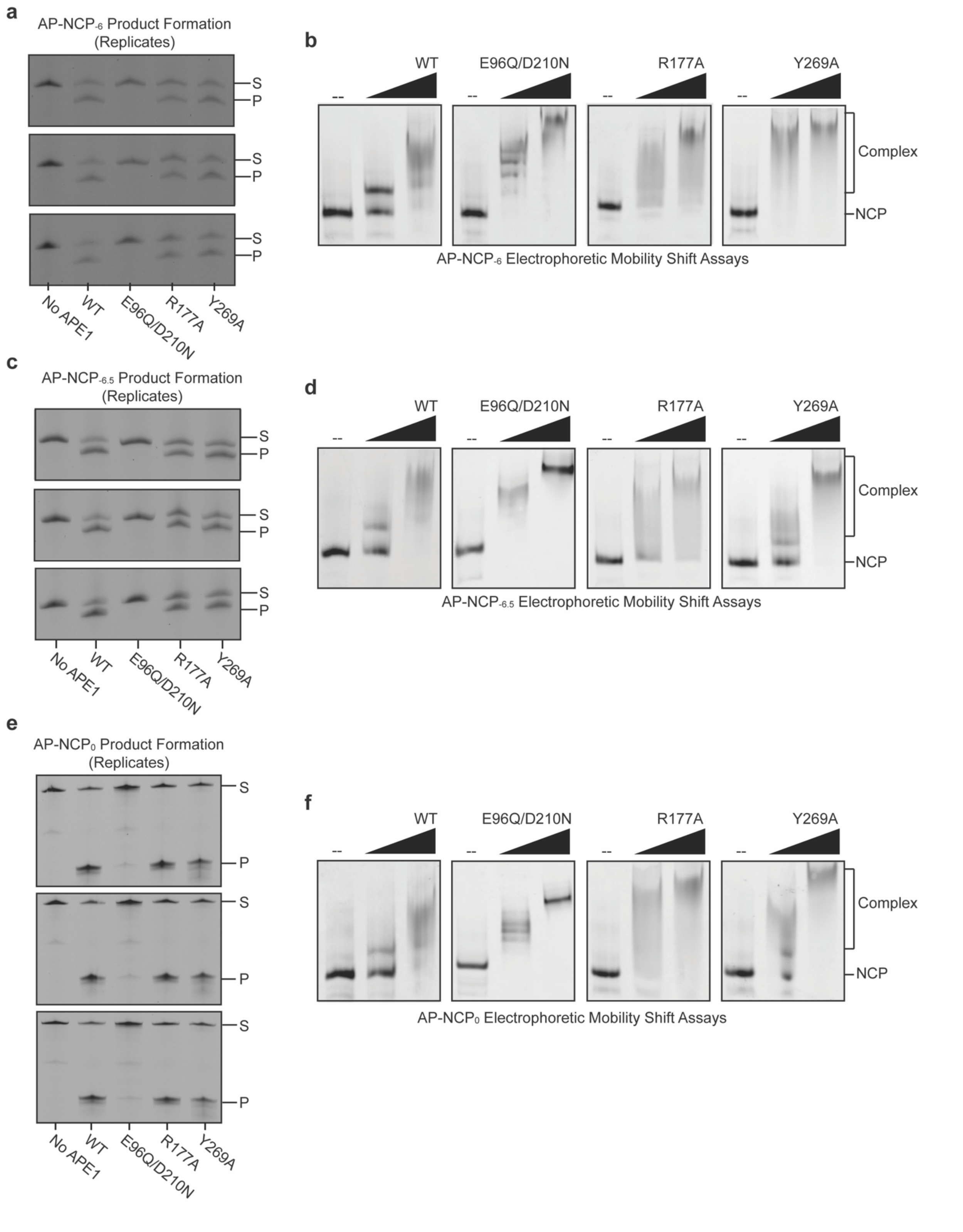

**Extended Data Fig. 5. Analysis of nucleosome binding and cleavage by APE1 mutants.**

**a,** Gels from three replicate AP-NCP$_{-6}$ product formation assays for WT, E96Q/D210N, R177A, and Y269A APE1. **b,** Representative gels of AP-NCP$_{-6}$ EMSA for WT, E96Q/D210N, R177A, and Y269A APE1. **c,** Gels from three replicate AP-NCP$_{-6.5}$ product formation assays for WT, E96Q/D210N, R177A, and Y269A APE1. **d,** Representative gels of AP-NCP$_{-6.5}$ EMSA for WT, E96Q/D210N, R177A, and Y269A APE1. **e,** Gels from three replicate AP-NCP$_{0}$ product formation assays for WT, E96Q/D210N, R177A, and Y269A APE1. **f,** Representative gels of AP-NCP$_{0}$ EMSA for WT, E96Q/D210N, R177A, and Y269A APE1. The substrate (S) and product (P) are labeled for all product formation assays.

### Extended Data Fig. 6. Single particle analysis of AP-NCP$_{-6.5}$.

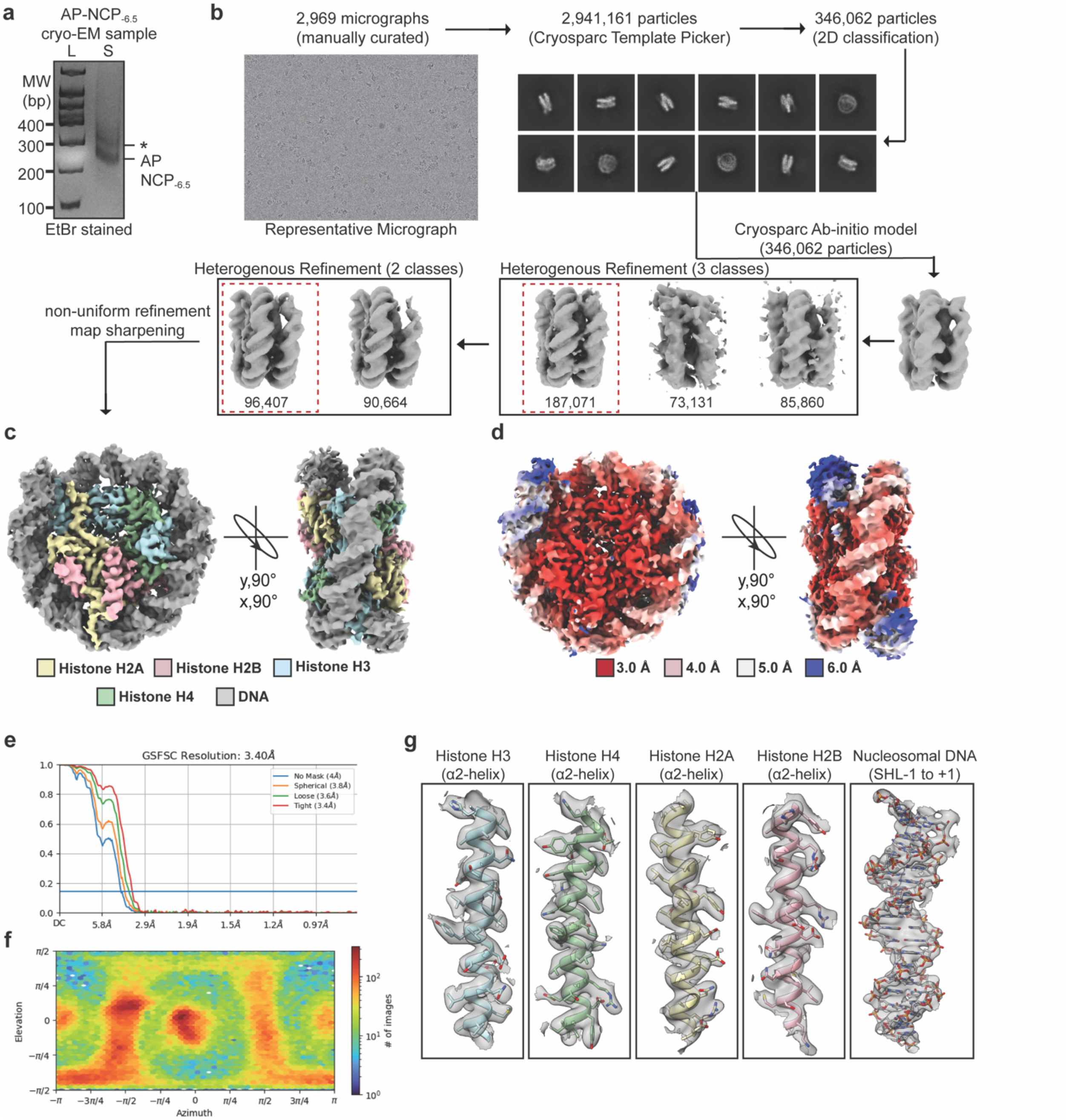

**Extended Data Fig. 6. Single particle analysis of AP-NCP$_{-6.5}$.**

**a,** Native PAGE gel of the purified AP-NCP$_{-6.5}$ cryo-EM sample. The 100 bp DNA ladder (L) and sample used to generate cryo-EM grids (S) are labeled. **b,** For single particle analysis, micrographs were manually curated and multiple rounds of 2D classification were performed yielding a final stack of 346,062 particles. A representative micrograph and set of 2D-classes are shown. An ab-initio model from 346,062 particles was generated before two rounds of heterogenous refinement (3 and 2 classes). All maps chosen for downstream analysis are labeled by a dotted red box. **c,** Final 3.4 Å sharpened cryo-EM map of AP-NCP$_{-6.5}$. **d,** Local resolution estimate for the AP-NCP$_{-6.5}$ cryo-EM map. **e,** Fourier shell correlation (FSC-0.143) for the AP-NCP$_{-6.5}$ map. **f,** Heatmap of the angular distribution of particles used to generate the final AP-NCP$_{-6.5}$ cryo-EM map. g) Representative segmented density for H2A, H2B, H3, H4 and the nucleosomal DNA from the AP-NCP$_{-6.5}$ cryo-EM map.

**Extended Data Fig. 7. Single particle analysis of AP-NCP$_0$.**

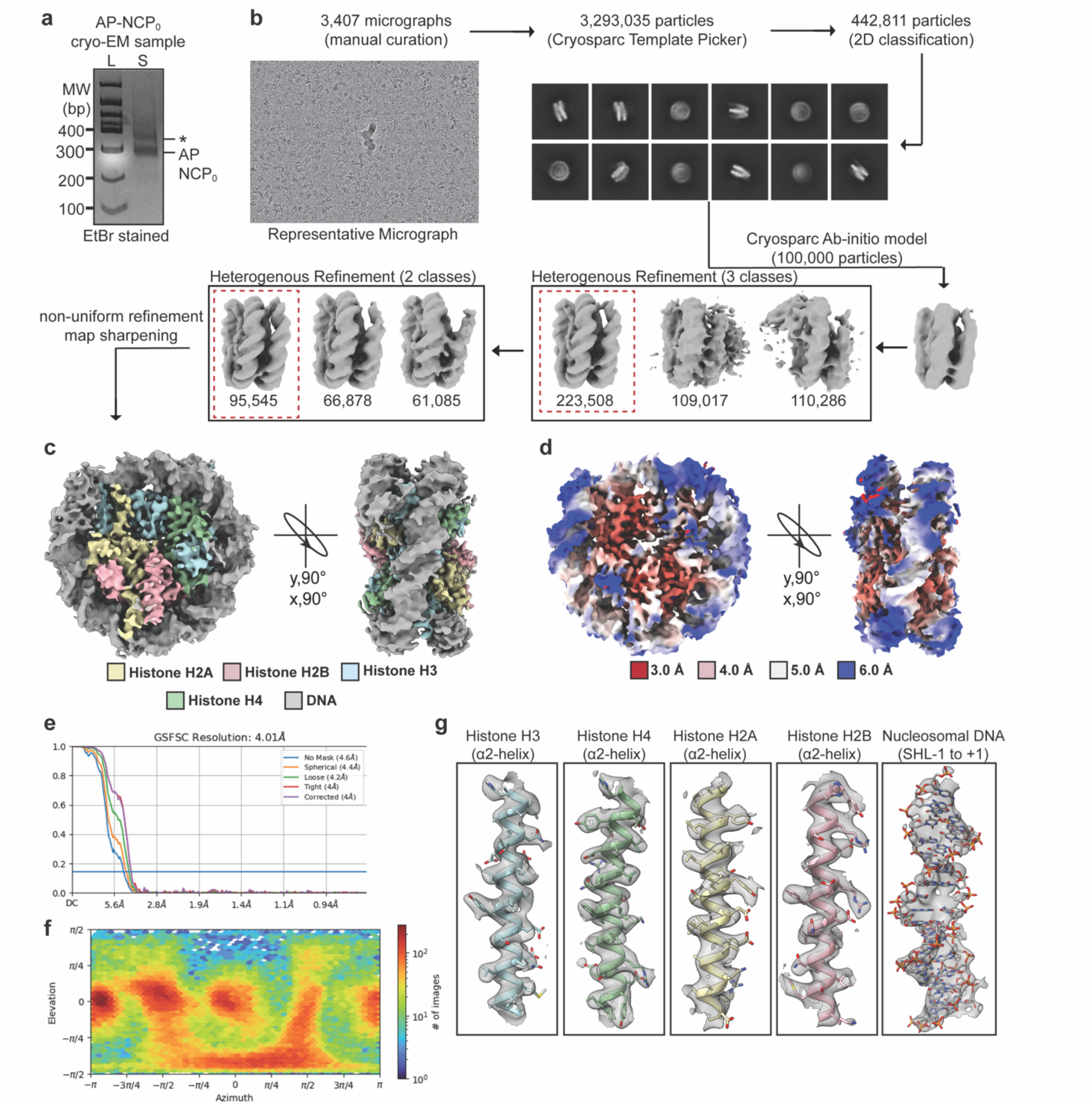

**Extended Data Fig. 7. Single particle analysis of AP-NCP$_0$.**

**a,** Native PAGE gel of the purified AP-NCP$_0$ cryo-EM sample. The 100 bp DNA ladder (L) and sample used to generate cryo-EM grids (S) are labeled. **b,** For single particle analysis, micrographs were manually curated and multiple rounds of 2D classification were performed yielding a final stack of 442,811 particles. A representative micrograph and set of 2D-classes are shown. An ab-initio model from 100,000 particles was generated before two rounds of heterogenous refinement (3 and 2 classes). All maps chosen for downstream analysis are labeled by a dotted red box. **c,** Final 4.0 Å sharpened cryo-EM map of AP-NCP$_0$. **d,** Local resolution estimate for the AP-NCP$_0$ cryo-EM map. **e,** Fourier shell correlation (cutoff-0.143) for the AP-NCP$_0$ map. **f,** Heatmap of the angular distribution of particles used to generate the final AP-NCP$_0$ cryo-EM map. g) Representative segmented density for H2A, H2B, H3, H4 and the nucleosomal DNA from the AP-NCP$_0$ cryo-EM map.

**Extended Data Fig. 8. Mechanism for APE1 processing occluded AP sites in the nucleosome.**

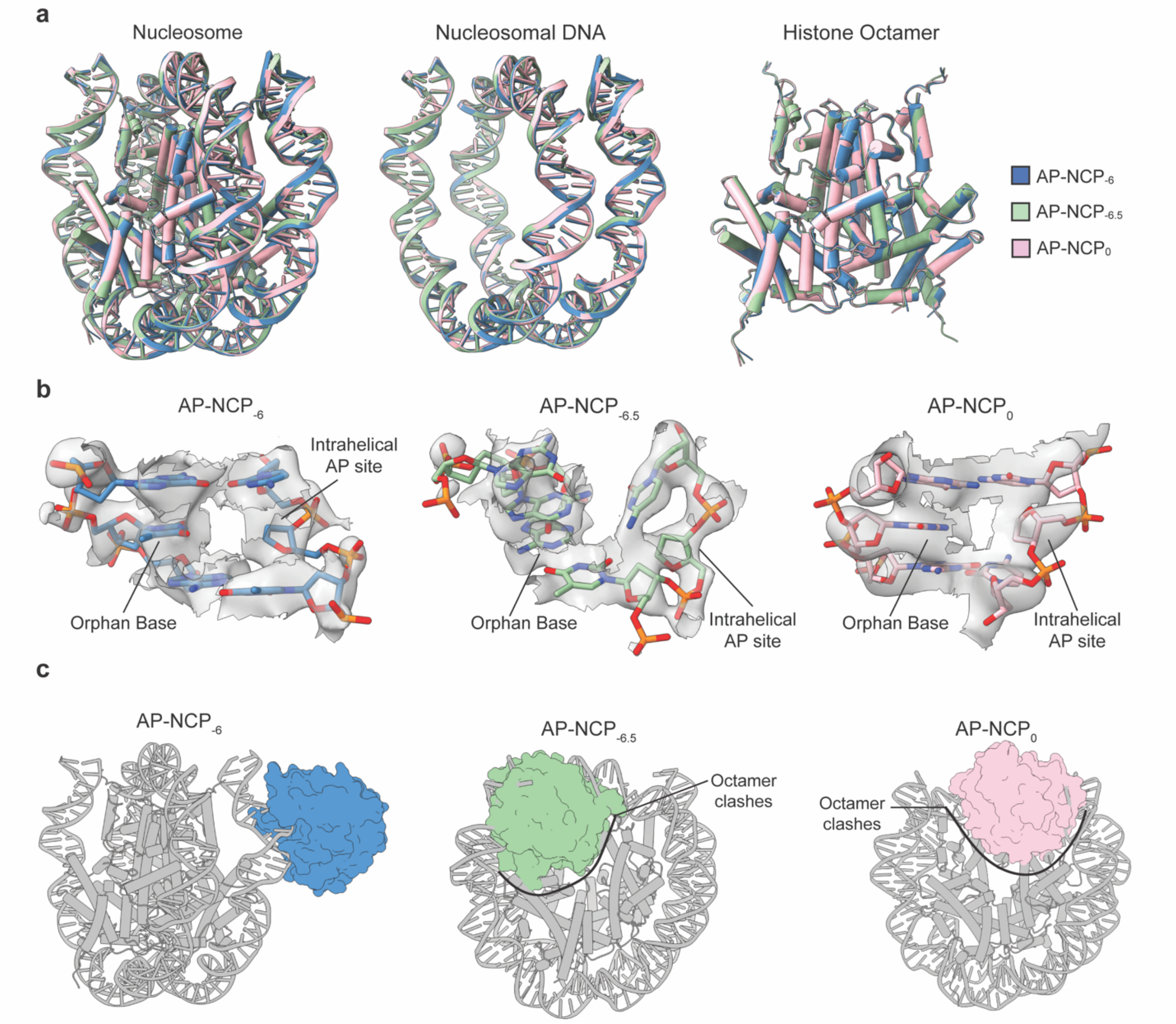

**Extended Data Fig. 8. Mechanism for APE1 processing occluded AP sites in the nucleosome.**

**a,** Overall comparison of the AP-NCP$_{-6}$, AP-NCP$_{-6.5}$, and AP-NCP$_0$ structures. **b,** Focused views of the intrahelical AP site for AP-NCP$_{-6}$ (left), AP-NCP$_{-6.5}$ (middle), and AP-NCP$_0$ (right). The cryo-EM map is shown as a transparent gray surface. **c,** Cryo-EM structure of the APE1-AP-NCP$_{-6}$ complex (left), and structural models of APE1 positioned to interact with the AP sites on AP-NCP$_{-6.5}$ (middle) and AP-NCP$_0$ (right). APE1 is shown in surface representation. Significant clashes of APE1 with the histone octamer are labeled.

**Extended Data table 1. Cryo-EM data collection, refinement, and validation.**

| **Structure** | **APE1-AP-NCP$_{-6}$** | **AP-NCP$_{-6}$** | **AP-NCP$_{-6.5}$** | **AP-NCP$_0$** |
|---|---|---|---|---|
| PDB accession | 7U50 | 7U51 | 7U52 | 7U53 |
| EMDB accession | EMD-26336 | EMD-26337 | EMD-26338 | EMD-26339 |
| **Data collection and processing** | | | | |
| Magnification | 29,000x | 29,000x | 130,000x | 29,000x |
| Voltage (kV) | 300 | 300 | 300 | 300 |
| Electron exposure ($e^-$/Å$^2$) | 50 | 50 | 47 | 50 |
| Defocus range (μm) | -0.8 to -2.2 | -0.8 to -2.2 | -0.7 to -2.1 | -0.8 to -2.0 |
| Pixel size (Å) | 0.40075 | 0.40075 | 0.415 | 0.40075 |
| Symmetry imposed | C1 | C1 | C1 | C1 |
| Initial particle images (no.) | 1,442,410 | 669,682 | 346,062 | 442,811 |
| Final particle images (no.) | 58,854 | 171,638 | 96,407 | 95,545 |
| Map resolution (Å) | 3.4 | 3.0 | 3.4 | 4.0 |
| FSC threshold | 0.143 | 0.143 | 0.143 | 0.143 |
| **Refinement** | | | | |
| Initial model used (PDB ID) | 4JJN, 5DFI | 4JJN | 4JJN | 4JJN |
| Model resolution (Å) | 3.4 | 3.0 | 3.3 | 3.9 |
| FSC threshold | 0.143 | 0.143 | 0.143 | 0.143 |
| **Model composition** | | | | |
| Nonhydrogen atoms | 14,035 | 11,883 | 11,847 | 11,828 |
| Protein residues | 1025 | 750 | 746 | 748 |
| Nucleotide | 288 | 290 | 290 | 288 |
| **B factors (Å$^2$ )** | | | | |
| Protein | 68 | 23 | 45 | 57 |
| Nucleotide | 158 | 85 | 118 | 131 |
| **r.m.s. deviations** | | | | |
| Bond Length (Å) (# > 4σ) | 0.009 (21) | 0.006 (9) | 0.005 (3) | 0.005 (5) |
| Bond Angles (°) (# > 4σ) | 1.189 (40) | 0.710 (8) | 0.663 (6) | 0.763 (24) |
| **Validation** | | | | |
| MolProbity score | 1.96 | 1.49 | 1.48 | 1.72 |
| Clashscore | 7.78 | 6.12 | 9.00 | 11.22 |
| Poor rotamers (%) | 2.45 | 0.80 | 0.48 | 1.29 |
| **Ramachandran plot** | | | | |
| Favored (%) | 96.4 | 97.1 | 98.8 | 97.7 |
| Allowed (%) | 3.6 | 2.9 | 1.2 | 2.3 |
| Disallowed (%) | 0 | 0 | 0 | 0 |

**Extended Data Table 2. Oligonucleotides for generating damaged nucleosomes.**

| Oligo | Sequence (5′ – 3′) |
|---|---|
| ND-NCP oligos | |
| ND_001 | ATCGGATGTATATATCTGACACGTGCCTGGAGACTAGGGAGTAATCCCCTTGGCGGTTAAAACGCGGGGGACAG |
| ND_002 | /5Phos/CGCGTACGTGCGTTTAAGCGGTGCTAGAGCTGTCTACGACCAATTGAGCGGCCTCGGCACCGGGATTCTCGAT |
| ND_003* | /56-FAM/ATCGAGAATCCCGGTGCCGAGGCCGCTCAATTGGTCGTAGACAGCTC |
| ND_004 | /5Phos/TAGCACCGCTTAAACGCACGTACGCGCTGTCCCCCGCGTTTTAACCGCCA |
| ND_005 | /5Phos/AGGGGATTACTCCCTAGTCTCCAGGCACGTGTCAGATATATACATCCGAT |
| AP-NCP$_{-6}$ oligos | |
| AP-NCP$_{-6}$_001 | ATCGGATGTATATATCTGACACGTGCCTGGAGACTAGGGAGTAATCCCCTTGGCGGTTAAAACGCGGGGGACAG |
| AP-NCP$_{-6}$_002 | /5Phos/CGCGTACGTGCGTTTAAGCGGTGCTAGAGCTGTCTACGACCAATTGAGCGGCCTCGGCACCGGGATTCTCGAT |
| AP-NCP$_{-6}$_003* | /56-FAM/ATCGAGAATCCCGGTGCCGAGGCCGCTCAATTGGTCGTAGACAGCTC |
| AP-NCP$_{-6}$_004 | /5Phos/TAGCACCGCTTAAACGCACGTACGCGCTGTCCCCCGCGTTTTAACCGCCA |
| AP-NCP$_{-6}$_005 | /5Phos/AGGGGATTACTCCCTAGTCTCCAGGCACGTGTCAGATATAT/idSp/CATCCGAT |
| AP-NCP$_{-6.5}$ oligos | |
| AP-NCP$_{-6.5}$_001 | ATCGGATGTATATATCTGACACGTGCCTGGAGACTAGGGAGTAATCCCCTTGGCGGTTAAAACGCGGGGGACAG |
| AP-NCP$_{-6.5}$_002 | /5Phos/CGCGTACGTGCGTTTAAGCGGTGCTAGAGCTGTCTACGACCAATTGAGCGGCCTCGGCACCGGGATTCTCGAT |
| AP-NCP$_{-6.5}$_003* | /56-FAM/ATCGAGAATCCCGGTGCCGAGGCCGCTCAATTGGTCGTAGACAGCTC |
| AP-NCP$_{-6.5}$_004 | /5Phos/TAGCACCGCTTAAACGCACGTACGCGCTGTCCCCCGCGTTTTAACCGCCA |
| AP-NCP$_{-6.5}$_005 | /5Phos/AGGGGATTACTCCCTAGTCTCCAGGCACGTGTCAGATATATACAT/idSp/CGAT |
| AP-NCP$_0$ oligos | |
| AP-NCP$_{-6.5}$_001 | ATCGGATGTATATATCTGACACGTGCCTGGAGACTAGGGAGTAATCCCCTTGGCGGTTAAAACGCGGGGGACAG |
| AP-NCP$_{-6.5}$_002 | /5Phos/CGCGTACGTGCGTTTAAGCGGTGCTAGAGCTGTCTACGACCAATTGAGCGGCCTCGGCACCGGGATTCTCGAT |
| AP-NCP$_{-6.5}$_003* | /56-FAM/ATCGAGAATCCCGGTGCCGAGGCCGCTCAATTGGTCGTAGACAGCTC |
| AP-NCP$_{-6.5}$_004 | /5Phos/TAGCACCGCTTAAACGCACGTACGC/idSp/CTGTCCCCCGCGTTTTAACCGCCA |
| AP-NCP$_{-6.5}$_005 | /5Phos/AGGGGATTACTCCCTAGTCTCCAGGCACGTGTCAGATATATACATCCGAT |

*These oligos do not contain 6-FAM label for cryo-EM substrates.
